# Transcriptional control of DNA repair networks by CDK7 regulates sensitivity to radiation in Myc-driven Medulloblastoma

**DOI:** 10.1101/2020.04.30.069237

**Authors:** Bethany Veo, Etienne Danis, Angela Pierce, Dong Wang, Susan Fosmire, Kelly D. Sullivan, Molishree Joshi, Santosh Khanal, Nathan Dahl, Sana Karam, Natalie Serkova, Sujatha Venkataraman, Rajeev Vibhakar

## Abstract

Myc-driven Medulloblastoma remains a major therapeutic challenge due to frequent metastasis and a poor 5-year survival rate. Myc gene amplification results in transcriptional dysregulation, proliferation, and survival of malignant cells. To identify therapeutic targets in Myc-amplified medulloblastoma we performed a CRISPR-Cas9 essentiality screen targeting 1140 genes annotated as the druggable genome. CDK7 was identified as a mediator of medulloblastoma tumorigenesis. Using covalent inhibitors and genetic depletion of CDK7 we observe the cessation of tumor growth in xenograft mouse models and increase in apoptotic mechanisms. The results are attributed to repression of a core set of Myc-driven transcriptional programs mediating DNA repair. We further establish that blocking CDK7 activity sensitizes cells to ionizing radiation leading to accrual of DNA damage and extended survival and tumor latency in medulloblastoma xenograft mouse models. Our studies establish a mechanism for selective inhibition of Myc-driven MB by CDK7 inhibition combined with radiation as a viable therapeutic strategy for Myc-amplified medulloblastoma.

## Introduction

Medulloblastoma (MB) is the most prevalent malignant brain tumor in children accounting for 15-20% of all pediatric brain tumors (Dhall, 2009). Therapeutic strategies for standard-risk patients consists of surgical resection followed by craniospinal irradiation and cytotoxic chemotherapy (Northcott et al., 2012; Ramaswamy et al., 2016). This strategy has improved patient outcomes for standard-risk patients, however high-risk patients still do poorly(Northcott et al., 2019; Ramaswamy et al., 2016). Previous molecular characterization of medulloblastoma identified tremendous heterogeneity and categorically subdivided MB into 4 main subgroups (*Wnt*, *Shh*, Group 3, and Group 4) with 12 subtypes(Kool et al., 2012; Schwalbe et al., 2017; Taylor et al., 2012). Group 3 patients and particularly Group 3γ, represent a severe form of the disease with a poor 5-year survival rate of less than 30%(Cavalli et al., 2017; Schwalbe et al., 2017). Group 3 tumors are associated with a higher incidence of metastatic disease, and less responsive to the standard therapeutic regimen often leading to tumor relapse. Many of these pathologies can be attributed to *c-myc* (*Myc*) gene amplification a main feature of group 3 MB(Ramaswamy et al., 2016). In tumor cells MYC functions as an oncogenic driver creating a dependency on transcriptionally active genes making MYC a master switch in controlling the transcription of the cancer genome(Bradner et al., 2017; Kress et al., 2015). Normally, MYC regulates essential cellular processes including proliferation, stem cell self-renewal, metabolism, translation, and ribosomal biogenesis(Adhikary and Eilers, 2005; van Riggelen et al., 2010). Pharmacologically targeting MYC remains challenging due to the widespread dependence of all cell types on these basic processes. Consequently, we sought to identify an alternative therapeutic target in MYC-driven medulloblastoma using an unbiased CRISPR-cas9 screen targeting the druggable genome. We identified the cyclin dependent kinase 7 as a top mediator of MB growth. CDK7 is a major transcriptional regulator responsible for facilitating RNA Pol II pre-initiation complex stability, pause, and release(Chen et al., 2018; Core and Adelman, 2019; Fisher, 2018; Glover-Cutter et al., 2009).

In this study, we investigate the mechanisms of CDK7 regulatory function in medulloblastoma growth and survival. We establish that genetic and chemical inhibition of CDK7 inhibits medulloblastoma proliferation and enhances apoptotic mechanisms in vitro and in xenograft PDX models. Mechanistically, we demonstrate that chemical inhibition of CDK7 disrupts RNA Pol II and MYC association with a core set of promoters mediating DNA repair. This action sensitizes MYC amplified MB cells response to irradiation lowering the effective dose of radiation for maximal cell damage. We further observe the combined radiation and CDK7 chemical inhibition amplifies tumor latency in a xenograft tumor model demonstrating the potential for pharmacological inhibition of CDK7 as a therapeutic strategy in MYC-amplified medulloblastoma.

## Results

### High expression of CDK7 in MYC-amplified medulloblastoma

In an effort to identify druggable targets in MYC-amplified medulloblastoma, we performed a CRISPR-cas9 screen targeting 1140 genes which at present includes genes for which there are current drugs/chemical inhibitors and genes that contain druggable pockets based on *in silico* analysis. Categories include kinases, epigenetic enzymes and metabolic enzymes among others (Figure S1). Utilizing three MYC-amplified medulloblastoma cell lines (D458, D425, and D341) we transduced cells and collected cell lysates before and 18 days after selection with puromycin (Figure 1A, 1B). DNA from the collected lysates were used for next-generation sequencing. The decrease in presence of strand guides indicates a decline in cell viability and therefore identifies genes essential to growth maintenance of MYC-amplifed MB (Figure 1A, 1C). Of the top strand guides found to be necessary for viability were genes that had previously been identified by us including *Wee1*, *Plk1*, *Brd4*, and *Ezh2* (Figure 1C) (Alimova et al., 2013; Alimova et al., 2012; Harris et al., 2014; Harris et al., 2012; Venkataraman et al., 2014). Within this group of genes, we found *Cdk7* to show a similar decrease in fold change when compared to before puromycin selected cells (Figure 1D). CDK7 (cyclin-dependent kinase) is a regulator of transcription initiation. As a part of the CDK-activating kinase (CAK) module in the TFIIH complex, CDK7 phosphorylates the Ser5 residue of RNA Pol II carboxy-terminal domain (CTD). The phosphorylation of Ser5 facilitates release from the pre-initiation complex and proximal pausing of RNA Pol II within 20 to 120 nucleotides of the TSS. These steps prime RNA Pol II for recruitment of the positive transcription elongation factor b (P-TEFb) which enables release of RNA Pol II into productive transcriptional elongation (Chen et al., 2018; Larochelle et al., 2007).

**Figure 1.**
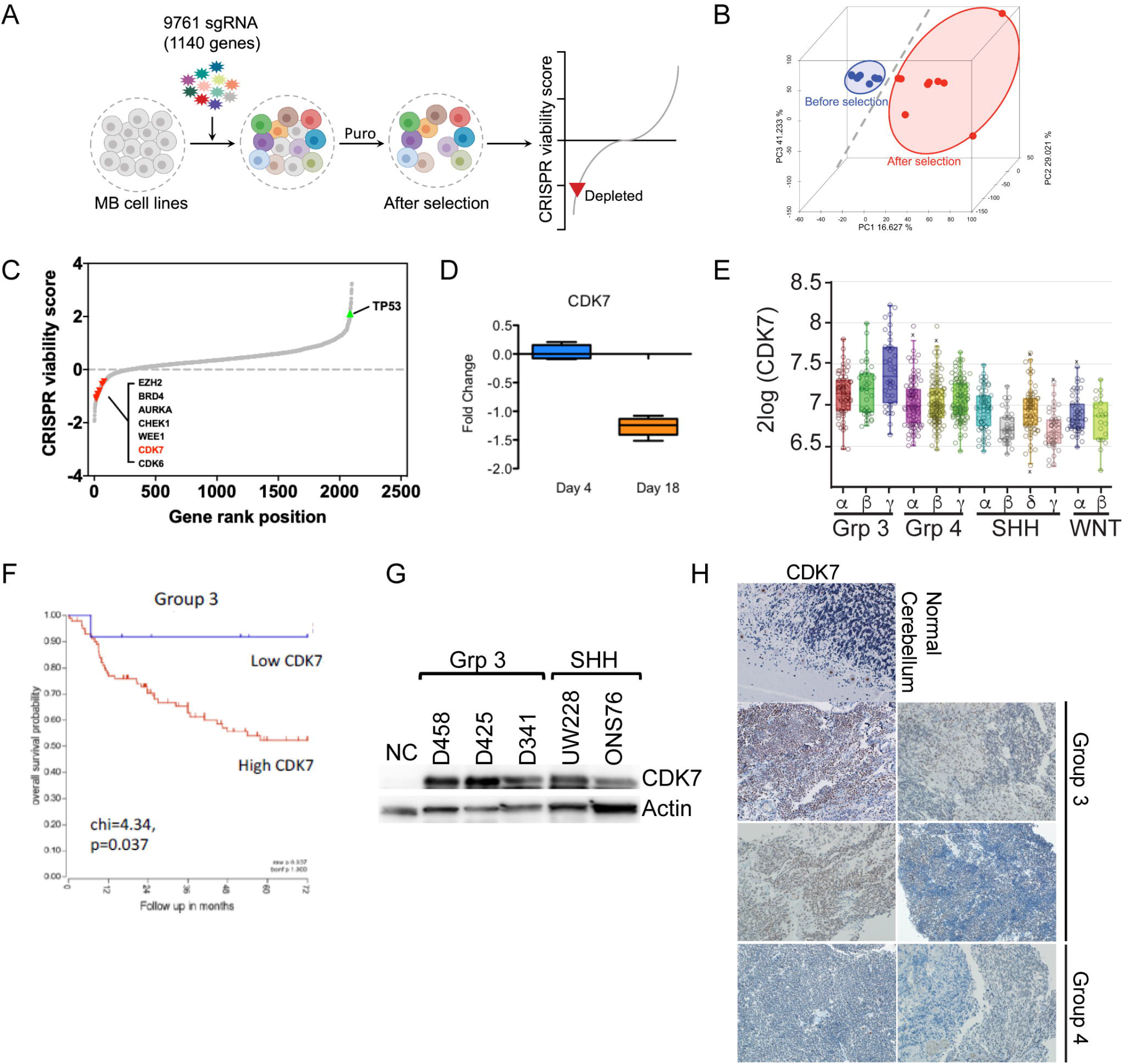
CDK7 expression in medulloblastoma enhanced in group 3 tumors. (A) Methodological graphic of the crispr-cas9 druggable kinase screen performed on three group 3 medulloblastoma cell lines. (B) PCA plot of before and after puromycin selection, also see S1 for heatmap. (C) S curve plot displaying the depletion of genes essential for Myc-MB cell line viability. Genes essential to Myc-MB growth previously identified by shRNA screen identified in black, new genes in red. (D) The expression of CDK7 before puromycin selection and 18 days after. (E) CDK7 expression in medulloblastoma by subtype. (F) Patient overall survival in Group 3 medulloblastoma in relation to CDK7 expression. (G) Immunoblot of CDK7 protein levels in three group 3 MB cell lines and two SHH MB cell lines. Quantification (S1) (H) Immunohistochemistry staining of CDK7 in normal cerebellum and 4 patient group 3 MB tumors and two group 4 tumors. Images shown at 20x.

To establish the relevant significance in patients, we investigated *Cdk7* gene expression across 763 patient tumor samples. *Cdk7* gene expression is associated with high levels of the *Myc* oncogene displaying the highest expression across group 3 and specifically group 3γ medulloblastoma (Figure 1E). Overall patient survival in group 3 medulloblastoma, shows a survival advantage in tumors with lower levels of *Cdk7* (Figure 1F). These results were also observed in our medulloblastoma cell lines with high (D458, D425), medium (DAOY, D283), and low (ONS76) levels of MYC (Figure 1G, S1). Using immunohistochemistry, we evaluated CDK7 protein levels in group 3 and group 4 tumors. Concordant with the mRNA gene expression data, we note a higher level of CDK7 in group 3 and group 4 tumor samples (Figure 1H). These results identify CDK7 as an important factor in the maintenance of medulloblastoma growth, and coordinate *Cdk7* expression with MYC abundance in the most aggressive form of medulloblastoma.

### Genetic knockdown of CDK7 represses medulloblastoma growth

To further determine the involvement of CDK7 in medulloblastoma proliferation, we performed genetic depletion studies by knocking down *Cdk7* with three independent shRNAs in MYC-amplified cell lines D458 and D425. Substantial reduction of CDK7 was achieved with all three shRNAs as shown by western blot (Figure 2A, S2). We evaluated phosphorylation of RNA Pol II at serine 5 and serine 2. Following depletion of CDK7, we verified that RNA Pol II Ser5 phosphorylation was decreased (Figure 2A, S2). Reduction of RNA Pol II Ser2 was also observed to a lesser degree than RNA Pol II Ser5. These results are in line with the role of CDK7 phosphorylation activating the catalytic subunit of P-TEFb (CDK9) which is responsible for phosphorylation of Ser2 (Glover-Cutter et al., 2009; Larochelle et al., 2007).

**Figure 2.**
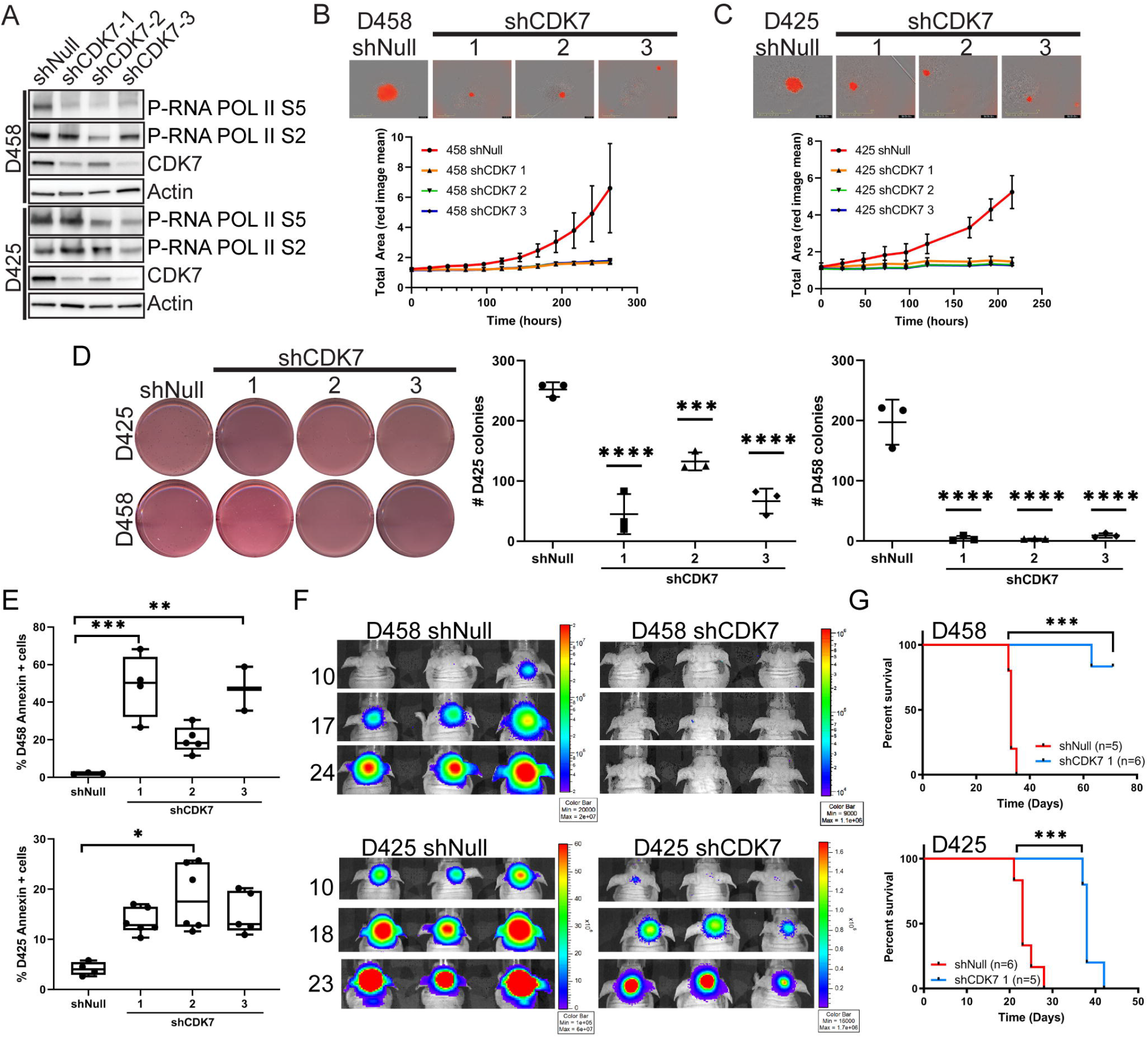
Genetic depletion of CDK7 decreases proliferation and tumor growth. (A) Immunoblot for indicated proteins of lysates transduced with three shRNAs against CDK7 or shNull. (B,C) Neurosphere assay growth in CDK7 depleted D458 or D425 MB cell lines. D458 or D425 cells expressing a NucRed™ marker were plated at 10 cells/well and monitored for growth using the Incucyte S3 over 10 days, representative images at 10 days are shown (above). Total average growth of neurospheres over 10 day period as determined by NucRed™ fluorescence (below). Experiment performed in triplicate, data shown as mean ± SD. (D) Methylcellulose assay of CDK7 knockdown cell lines (D458,D425; shown left). Colony count for D425 (middle) and D458 (right). Experiment performed in triplicate. Shown is a scatter plot with mean and ±SD. Statistical analysis, two-way Anova. ****,p<0.0001; ***,p<0.001. (E) Annexin V (+) staining assay. D458 (top) and D425 (bottom) knockdown cells were stained for Annexin V and measured by guava flow cytometry. Box and whisker plot of the % (+) Annexin V stained cells (y-axis) is shown. Experiments performed in triplicate. Statistical analysis, two-way nova. ***,p<0.001; **,p<0.005; *,p<0.05. (F) Representative bioluminescence images of xenograft D458 CDK7 knockdown or shNull (top) and D425 shCDK7 or shNull (bottom). Days post injection indicated on the left. Color scales indicate bioluminescence radiance in photons/sec/cm^2^/steradian shown right. (G) Kaplan-Meier survival curve of D458 CDK7 knockdown (n=6) and shNull (n=5), (top). D425 shNull (n=6) or shCDK7 (n=5) (bottom). Statistical analysis log-rank (Mantel-Cox) test; ***,p<0.001.

Medulloblastoma is thought to include brain tumor stem cells which promote proliferation, cancer stem cell self-renewal, and resistance to traditional therapeutic interventions (Singh et al., 2003). Following knockdown of CDK7 we assessed cancer stem cell self-renewal and proliferation through a neurosphere growth assay in D458 and D425 cell lines. Expression of the NucRed marker in D458 and D425 shNull and CDK7 KD cell lines were monitored on the Incucyte S3 system over 10 days. CDK7 depletion shows substantial inhibition of neurosphere growth as compared to the shNull in both D458 and D425 cells (Figure 2B, C). To further understand if neurospheres failed to grow initially or if self-renewal was disrupted, 10-day old neurospheres were collected and re-plated as secondary spheres for another 10 days. Similarly, CDK7 depleted cells were unable to sustain growth, whereas shNull D458 and D425 neurospheres continued to grow and develop new neurospheres (Figure S2). This data suggests that loss of CDK7 prevents further proliferation of medulloblastoma cells and interrupts the self-renewal of MB cancer stem cells. To then establish whether CDK7 loss impaired the ability of MB cells to form adhesion independent colonies we performed a methylcellulose assay. ShCDK7 transduced D458 and D425 cells were plated in 1.3% methylcellulose and colonies counted after 10 days. CDK7 depleted cell lines showed a 50% reduction in the number of colonies in D425 cells compared with controls, whereas in D458 cells fewer than 50 colonies formed (Figure 2D). To determine whether the observed decrease in proliferation was due to a cessation of growth or an increase in cell death, we examined Annexin V positivity by flow cytometry. D458 cells depleted of CDK7 showed a 40-60% increase in Annexin V (+) suggesting that loss of CDK7 leads to an increase in apoptotic cells (Figure 2E). Likewise, D425 shCDK7 cells increased Annexin V (+) by 15-25%. These results show CDK7 is an essential component to MYC-amplified medulloblastoma growth, and reduction of CDK7 disrupts MB propagation and self-renewal by enhancing apoptosis.

To evaluate whether CDK7 depletion effects tumor growth in vivo, we utilized bioluminescent CDK7 knockdown D458 and D425 cells for cerebellar injections into nude mice. Mice were monitored for tumor growth with periodic bioluminescence imaging to evaluate tumor burden. Control mice were sacrificed due to tumor burden by 32 days post injection (Figure 2F). In contrast, shCDK7 D458 injected mice had not formed visible tumors as detected by bioluminescence at 24 days post-injection (Figure 2F). Imaging of D458 shCDK7 mice was continued up to 75 days post-injection (Figure S2). At end-point, a single mouse had succumbed to tumor burden by day 62 whereas all others remained healthy (Figure 2G). Similarly, D425 shNull mice developed tumors by day 10 and needed to be sacrificed by day 25 due to tumor burden. Delayed tumor formation was observed in 425 shCDK7 injected mice where tumors were evident by day 18 (Figure 2F). Survival in D425 shCDK7 mice was enhanced by 10 days compared to shNull injected mice (Figure 2G). These in vivo results show reducing *Cdk7* expression dramatically increased the time to tumor burden and in the case of D458 cells prevented tumor growth all together in 5 of 6 mice. D425 cells carry a *p53* mutation in addition to *Myc* amplification which may impact tumor survival. Taken together, the genetic depletion of *Cdk7* confirms the results from the CRISPR-cas9 druggable screen and establishes CDK7 as an essential factor mediating MYC-amplified medulloblastoma growth.

### CDK7 chemical inhibition displays potent selectivity to MYC-amplified medulloblastoma

Several covalent inhibitors of CDK7 display promise as clinically relevant inhibitors and are currently in phase I clinical trial for advanced solid tumors (SCLC, prostate, ovarian, and TNBC). Among them THZ1 and THZ2 show selectivity for CDK7 at low nanomolar doses, though it has been reported to target CDK12/13 at higher concentrations of >150nM (Kwiatkowski et al., 2014). Additionally, THZ2 derived from THZ1 maintains the specificity of THZ1 while increasing the half-life of the compound (Wang et al., 2015). To determine if group 3 medulloblastoma cell lines were susceptible to THZ1 and THZ2 inhibition we treated Myc-amplified cell lines D458 and D425 in addition to non-MYC amplified cell lines ONS76 and UW228. In MYC-amplified cells D458 and D425, THZ1 showed considerable potency with an IC_50_ range of 10nM-13nM (Figure 3A). Whereas UW228 and ONS76 cell lines, which represent SHH type tumors, displayed a 10- and 20-fold increase in IC_50_ at 140 and 240nM respectively (Figure 3A). Similar to THZ1, THZ2 was more potent in D458 and D425 cells (Figure 3A). These results suggest that effective CDK7 inhibition requires MYC-amplification in medulloblastoma. Additional tumors which exhibit MYC transcriptional addiction are reported to respond to THZ1 albeit at higher concentrations than we have found (Christensen et al., 2014; Lu et al., 2019; Nagaraja et al., 2017; Wang et al., 2015).

**Figure 3.**
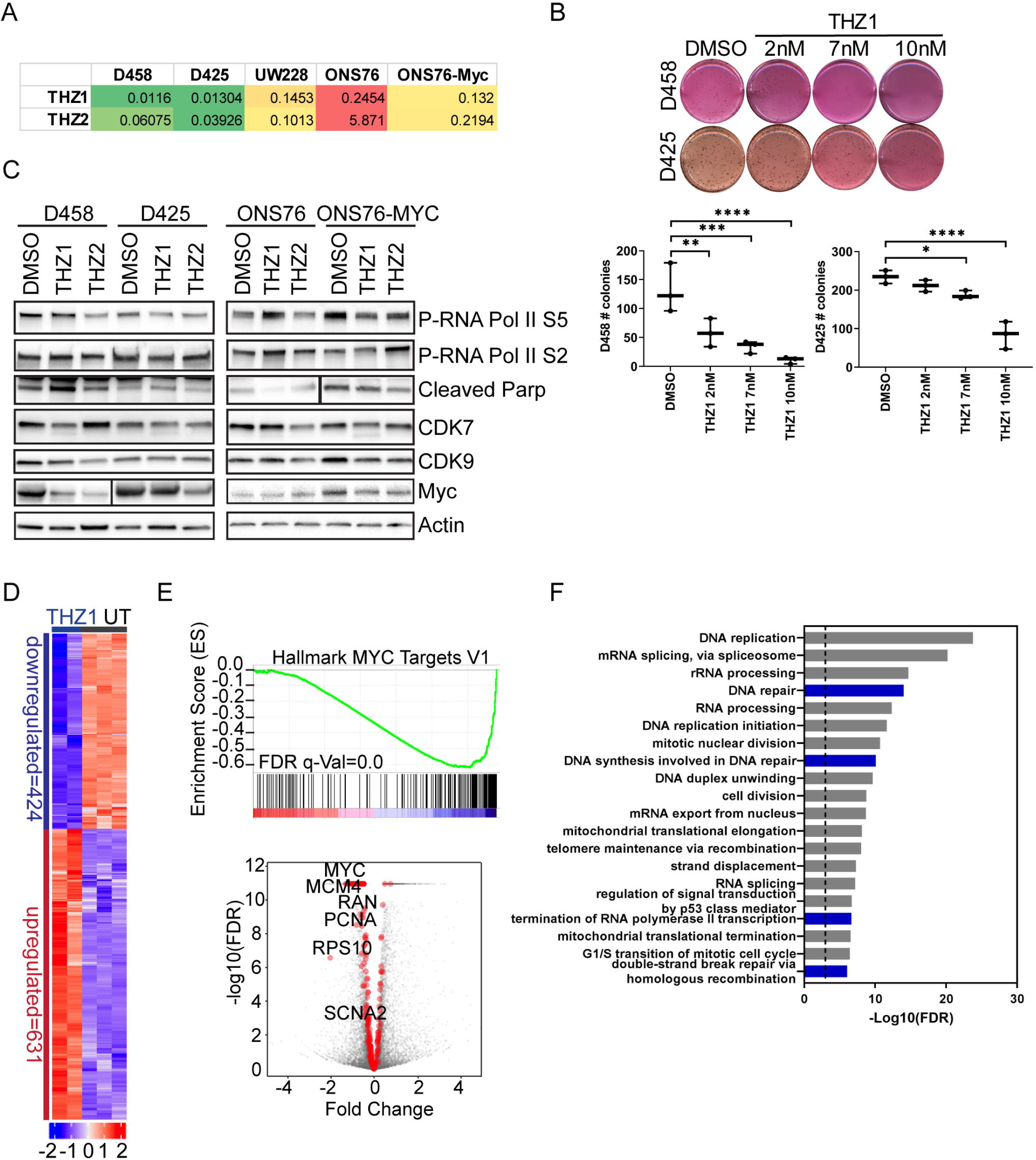
Low dose THZ1 silences Myc activated genes. (A) Log IC50 determination of THZ1 and THZ2 on group 3 MB cell lines and non-MYC amplified cell lines. Shown is IC50 in μM concentrations. Color scale indicates low (green) to high (red) concentrations.(B) Representative images of methylcellulose assay of D458 or D425 cells treated with DMSO, 2 (IC20), 7 (IC30), 10nM (IC50) THZ1, (top). Scatter plot mean ± SD indicating the number of colonies/well for D458 and D425 (bottom). Statistical analysis one-way Anova. ****,p<0.0001;***,p<0.001; **,p<0.01. (C) Immunoblot of proteins from D458, D425, ONS76, and ONS76-MYC treated with THZ1 or THZ2 IC50 for 48hrs. (D) RNA-seq gene expression heat map of THZ1 IC50 (10nM) treated D458 for 48hrs and untreated cells. Calculated Z score scale shown. (E) Gene set enrichment analysis Hallmark MYC targets V1 of genes downregulated by THZ1 (10nM) treatment. FDR q-Value=0.0. (top). Volcano plot of gene expression changes in hallmark MYC targets V2, shown in red. All genes in gray. (bottom). (F) GO terms for THZ1 10nM treated D458 of downregulated gene set from RNA-seq data analyzed using DAVID. Shown is −Log10(FDR) values where the dotted line indicates FDR<0.001. DNA repair related categories shown in blue.

In order to determine if the MYC-amplified MB cells response to THZ1 recapitulates genetic depletion of *Cdk7* we first examined MB cell propagation as neurospheres. D458 NucRed cells were seeded as neurospheres and monitored over 10 days for growth in the presence of increasing concentrations of THZ1. Higher concentrations ranging from 2μM-50 nM completely inhibited neurosphere growth over the 10-day period, whereas minimal growth was observed at 10nM THZ1 (Figure S2). Compared with DMSO, the striking inhibition of neurosphere proliferation with THZ1 mimics the genetic KD of *Cdk7* observed previously. We then evaluated independent colony formation of D458 and D425 cell lines exposed to THZ1 at IC_20_, IC_30_, and IC_50_. A limited number of colonies formed at 2nM in the D458 cell line, however 7 and 10 nM concentrations significantly reduced colonies from both D458 and D425 cells (Figure 3B). These results recapitulate our genetic KD study and show that THZ1 effectively inhibits independent colony growth as well as neurosphere growth.

We then evaluated the dependency of THZ1 on MYC-amplification and addressed the selectivity of THZ1 and THZ2 for CDK7. An inducible *Myc* overexpression ONS76 cell line was used to discern whether the presence of MYC enhanced the response to CDK7 chemical inhibition in comparison to wild type ONS76 cells. High *Myc* expressing MB cells (D458 and D425) as well as ONS76 and ONS76-MYC were exposed to 50nM THZ1 and THZ2 for 48hrs, to elicit a defined response. We examined Ser5 phosphorylation on RNA Pol II as a read out of CDK7 inhibition. Treatment reduced Ser5 phosphorylation in D458 and D425 cells in addition to the ONS76 cell lines (Figure 3C, S2). RNA Pol II Ser2 phosphorylation is attributed to CDK12 and CDK9 kinase activity, however treatment does not appear to effect Ser2 phosphorylation in all cell lines indicating the selectivity of THZ1 and THZ2 (Figure 3C, S2). Additionally, we blotted for CDK7 and CDK9 but observed little change with treatment (Figure 3C, S2).

To evaluate if treatment with THZ1 and THZ2 induces cell death we looked for cleavage of PARP. In D458 and D425 cells we observed an increase in cleaved PARP (Figure 3C, S2). There was minimal cleavage of PARP in ONS76 cells. However, upon induction of *Myc* in ONS76 cells there was significant enhancement of cleaved PARP with treatment (Figure 3C, S2). These results suggest THZ1 and THZ2 induce apoptosis and that enhancement of apoptosis may be reliant on the presence of MYC. Since THZ1 and THZ2 treatment disrupts MYC-transcriptional dependencies, we examined if MYC protein levels were altered by treatment. In D458 and D425 cells, a reduced level of MYC is observed with THZ1 and is more prominent with THZ2 treatment (Figure 3C, S2). Though the ONS76 cell line has lower levels of MYC protein we did not find any change with THZ1 or THZ2 treatment. However, under *Myc*-inducible conditions an increase in MYC is observed which is then reversed by THZ1 and THZ2 (Figure 3C, S2). These results demonstrate THZ1 and THZ2 selectively prevents CDK7 catalytic activity limiting RNA Pol II Ser5 phosphorylation. The reduction in Ser5 phosphorylation holds RNA Pol II in the pre-initiation complex preventing RNA Pol II release into proximal promoter pausing and eventual productive elongation thus reducing transcription of highly active genes. Consistent with this, MYC protein levels decrease with THZ1 and THZ2 treatment, disrupting MYC transcriptional dependency, and enhancing apoptotic mechanisms.

To fully understand the transcriptional response to THZ1 treatment in group 3 medulloblastoma cells, we performed a global gene expression analysis on D458 cells treated with THZ1 at IC_50_ for 48 hrs. Significant differential gene expression was set at P<0.05 and shows 40% of genes decreased expression (Figure 3D). Gene set enrichment analysis broadly shows the depletion of MYC transcriptional targets (Figure 3E). Gene ontology (GO) analysis performed on the downregulated genes show a significant association with DNA repair, double stranded break repair, termination of transcription, and replication (Figure 3F). Global gene expression analysis indicates that even at 10nM THZ1 group 3 medulloblastoma cells respond dramatically to CDK7 inhibition by abrogating the effect of MYC and inducing cell death mechanisms.

### CDK7 inhibitor effects in orthotopic and patient-derived xenograft mouse models of medulloblastoma

We showed that THZ1 and THZ2 have selectivity and efficacy against MYC-amplified medulloblastoma cell lines. To determine the relative potency of CDK7 chemical inhibition in vivo we tested THZ2 treatment in orthotopic mouse models of medulloblastoma growth. We elected to use THZ2 due to the enhanced half-life of THZ2 in vivo (Wang et al., 2015). Bioluminescent D458 or D425 cells were injected into murine cerebellum to generate orthotopic xenografts. Mice were then randomized and tumor growth was verified by IVIS imaging by day 10 post-injection. Intraperitoneal injections of THZ2 at 15mg/kg or vehicle were then conducted daily for 25 days. D458 orthotopic mice exposed to vehicle displayed tumor growth consistent with our previous reports showing the first signs of tumor ranging from day 10-12 (Figure 4A) (Veo et al., 2019). Initial IVIS signals showed similar tumor sizes among all cohorts, however tumor growth continued to slow in THZ2 treated mice significantly reducing the average IVIS signal (Figure 4B). All vehicle treated mice had extensive tumor burden 30 days post-injection, and had to be euthanized. Whereas 5 of the 6 D458 THZ2 treated cohort were still alive at the same time point (Figure 4F). THZ2 treated D458 mice survived on average 2.5 days longer compared to vehicle treated mice (Figure 4F). Similarly, D425 orthotopic mice treated with vehicle displayed progressive tumor growth (Figure 4C, D). The D425 cell line grows more aggressively, and initial tumors were first identified by IVIS on day 8 post-injection. Where 7 of 9 vehicle-treated mice succumbed to tumor burden and euthanasia by day 28 (Figure 4F). The D425 THZ2 treated cohort displayed slowed progression of tumor growth by IVIS with 5 of 9 still alive at the same time point (Figure 4D, F). Survival in D425 THZ2 treated mice was extended by 3 days (Figure 4F).

**Figure 4.**
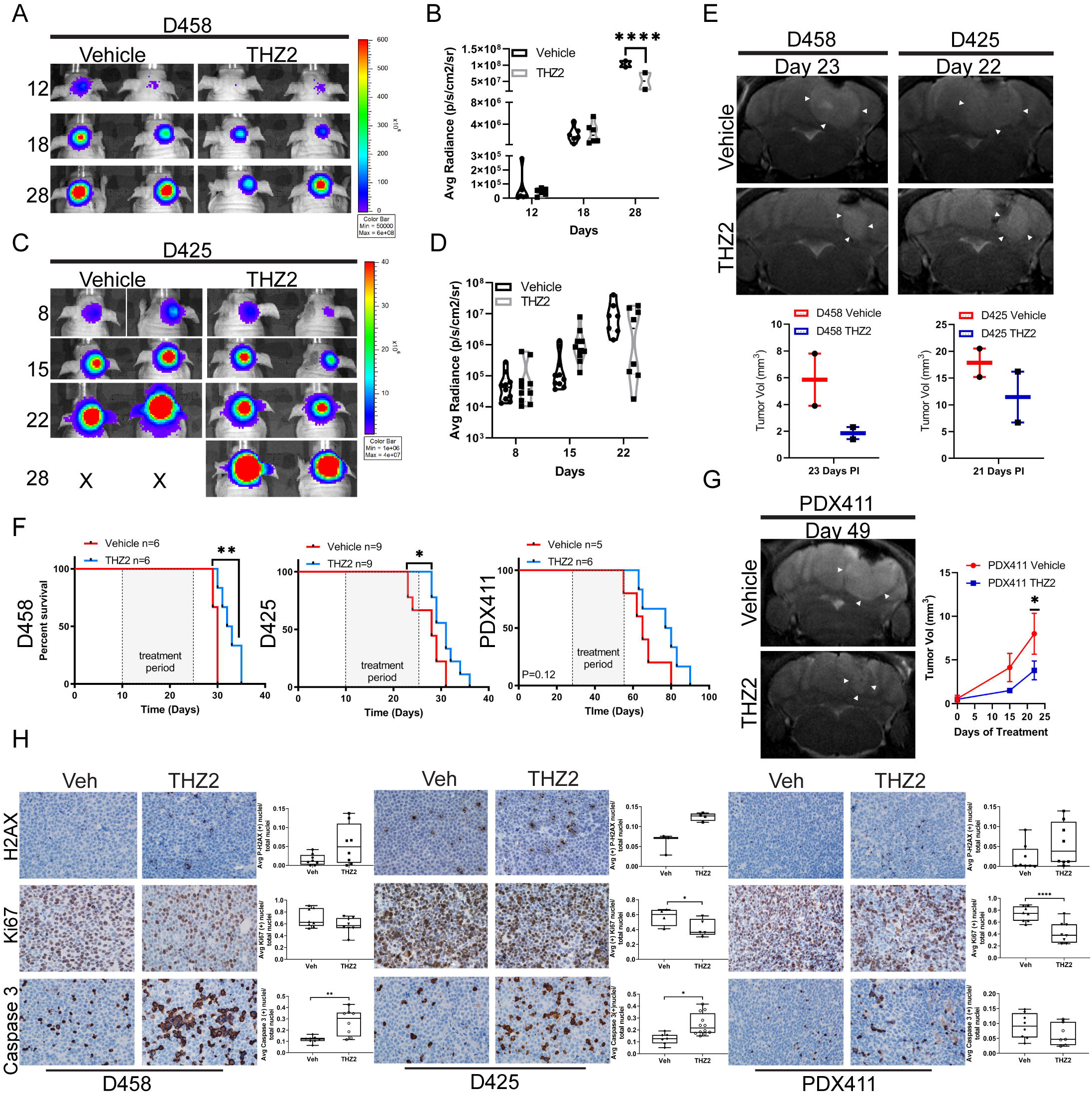
In vivo efficacy of CDK7 inhibition. (A) Nude mice xenografts injected with D458 cells were treated with vehicle (n=6) or THZ2 (n=6) 15mg/kg daily. Representative images of bioluminescence imaging conducted at 12, 18, and 28 days post-injection is shown. Color scales indicate bioluminescence radiance in photons/sec/cm^2^/steradian shown right. (B) Violin plot of mean total radiance in photons/sec/cm^2^/steradian. Statistical analysis two-way Anova, ****, p<0.0001. (C) D425 nude mice xenografts treated with vehicle (n=9) and THZ2 (n=9) 15mg/kg daily. Representative images show bioluminescence imaging at days 8, 15, and 22 post-injection. Color scales indicate bioluminescence radiance in photons/sec/cm^2^/steradian shown right. X indicates mouse death. (D) Violin plot of mean total radiance in photons/sec/cm^2^/steradian. (E) Representative MRI of vehicle and THZ2 treated D458 and D425 xenografts (top). White arrows indicate tumor edge. (Bottom) Tumor volume in mm^3^ (y-axis) in vehicle and THZ2 treated mice for D458 (left) and D425 (right). (F) Kaplan-Meier survival curve of D458, D425, and PDX411 xenograft mice treated with vehicle or THZ2. Treatment period for 25 days indicated in shaded box. Statistical analysis log-rank (Mantel-Cox) test, **,p<0.005; *,p<0.05. (G) PDX411 MRI and corresponding tumor volume in mm^3^ (y-axis) at 28, 42, and 49 days post-injection. Statistical analysis two-way Anova, *,p<0.05. (H) Representative immunohistochemistry staining for P-H2AX, Ki67, and caspase 3 of D458, D425, and PDX411 tumors from vehicle and THZ2 treated mice. Images taken at 40x. Statistical analysis one-way Anova. ****,p<0.0001; **,p<0.01; *p<0.05.

MRIs were performed on vehicle and THZ2 treated cohorts from D458 and D425 mice to examine tumor volumes and brain morphology. The average tumor volume from THZ2 treated cohorts was smaller in both models, agreeing with IVIS imaging (Figure 4E). Additionally, vehicle-treated cohorts exhibited edema and tumor invasiveness into the ventricular space one week earlier compared with THZ2 treated mice (Figure S3).

The efficacy of THZ2 in medulloblastoma cell line orthotopic xenografts instigated more direct studies using the patient-derived xenograft, MED411FH (PDX411) (Cook Sangar et al., 2017). MED411FH was originally harvested from a large cell anaplastic medulloblastoma that was molecularly characterized as a group 3 tumor. To enhance our probability of tumor uptake, NSG mice were used rather than nude mice which, in our hands, display a higher take percentage compared to the nude mouse. Following intracranial injections of MED411FH into the cerebella of NSG mice, mice were randomized. At 28 days post injection MRI confirmed the presence of tumors in three representative mice. Mice began treatment with THZ2 at 15mg/kg daily for 25 days with bi-monthly MRIs to monitor tumor size. MRIs of the vehicle treated mice displayed early signs of edema and visible tumor in the ventricular space at 42 days (Figure S3). In contrast, within the first 15 days of THZ2 treatment PDX411 mice displayed reduced tumor volumes by MRI (Figure 4G). Tumor progression maintained a slower overall course in THZ2 treated cohort with 4 of 6 mice still alive on day 68, whereas 4 of 5 vehicle-treated mice had succumbed to tumor burden and were euthanized. The THZ2 treated cohort survived on average 13.5 days longer than the vehicle-treated cohort (Figure 4F).

Immunohistochemistry was performed on collected mouse tissue at endpoint from each individual cohort. Initially, Ki67 staining was used to examine tumor proliferation in vivo. In THZ2 treated tumors, Ki67 staining reduced by 31% in PDX411 tumors (P<0.0001) and by 15 % in D425 tumors (P<0.05) compared to vehicle treated tumors (Figure 4H). Global gene expression analysis identified DNA repair pathways as being downregulated by THZ1 and THZ2 treatment.

To determine if THZ2 increased DNA damage within the tumors P-γH2AX staining was done. THZ2 treated tumors increased P-γH2AX staining by 12% in D425 tumors and trended upwards in D458 and PDX411 tumors compared to vehicle treated tumors (Figure 4H). Importantly DNA damage was restrained to tumors and not found in regions of sustained neurogenesis, specifically the hippocampus (Figure S3). We then evaluated Cleaved caspase 3 staining to identify apoptotic tumor cells. THZ2 treated mice increased Cleaved caspase 3 staining by 16% in D458 tumors (P<0.01) and 12% in D425 tumors (P<0.05) (Figure 4H). Likewise, activated caspase 3 staining was not found in the resounding brain or hippocampus (Figure S3).

Our data suggests THZ2 is a specific potent inhibitor of CDK7, though the presence of unmanageable chemical toxicity can negate the positive effects of a drug. To be sure that chemical toxicity did not produce adverse side effects in mice, blood samples were collected from mice at the end of experiment. Measuring counts of WBC, lymphocytes, and neutrophils showed no significant differences between vehicle and THZ2 treated mice in D458 and PDX411 (Figure S3). Similarly, platelets and hemoglobin counts indicated no significant changes with THZ2 treatment compared with vehicle in D458 and PDX411 (Figure S3). D425 mouse experiments were completed prior to beginning CBC collections. These results indicate THZ2 inhibition of CDK7 reduces tumor proliferation and increased apoptosis in vivo, while minimizing adverse neurological side effects.

### CDK7 inhibition decreases RNA Pol II and MYC promoter occupancy in a subset of genes

CDK7 inhibition effects transcriptional regulation through phosphorylation of RNA Pol II CTD at Ser5. To examine the mechanisms of CDK7 activity in MYC MB we evaluated MYC and RNA Pol II promoter occupancy with and without CDK7 inhibition. We performed ChIP sequencing analysis using RNA Pol II and MYC antibodies in both D458 and D425 cell lines that were treated with 10nM THZ1. We found the disruption of CDK7 activity prevents RNA Pol II promoter loading near the transcriptional start site in a subset of genes (Figure 5A). We identified 3 gene clusters of RNA Pol II occupancy. Cluster 3 demonstrates genes not transcriptionally active. Importantly, sweeping decreases in global Pol II poised transcription were not observed. Cluster 2 indicates retention of Pol II binding on genes mediating mRNA processing and translation as well as ribosome biogenesis (Figure 5A and S4). Cluster 1 displays the most dramatic decrease in RNA Pol II occupancy at the TSS. Gene ontology analysis identified genes primarily associated with DNA repair, DNA replication, and ncRNA processing (Figure 5A). ChIP-sequencing of MYC shows decreased association of MYC at the TSS with CDK7 inhibition, in line with our transcriptomic analysis (Figure 5C). Clustering of MYC ChIP profiles identified select areas of decreased occupancy at the TSS rather than a blanketed decrease in promoter association (Figure 5C). Cluster 2 and cluster 3 exhibit little change in MYC occupancy displaying regions of continued MYC transcriptional activity including translation and RNA catabolic processing, and regions of transcriptional inactivity (Figure 5C and S4). Depleted MYC promoter association was most prominent in cluster 1, with DNA repair processes topping the functional ontology list (Figure 5C). To discern whether decreased promoter occupancy of RNA Pol II and MYC overlapped onto genes downregulated from the transcriptomic analyses we compared the first cluster from RNA Pol II and MYC ChIP profiles with downregulated genes. Of the genes downregulated by THZ1, 122 genes matched with promoter sites that also had decreasing RNA Pol II and MYC indicating that not all transcriptionally inhibited genes were MYC targets and not all MYC targets were transcriptionally repressed by CDK7 inhibition (Figure 5E). Functional annotation of the overlapping genes indicates a significant association with DNA repair, replication, and homologous recombination (HR) processes (Figure S4). Further examination of the top genes in the DNA repair pathway of cluster 1 show the association of RNA pol II and MYC was significantly decreased at the TSS of genes mediating HR including *Brca2*, *Brcc3*, *Rad51c*, and *Xrcc2* (Figure 5F). Whereas the top genes in mRNA processing of cluster 2 have similar overall occupancy for both RNA Pol II and MYC (Figure 5F). These results emphasize that inhibition of CDK7 activity diminishes RNA Pol II and MYC promoter occupancy at specific gene clusters. Further, the functional overlap between RNA Pol II and MYC produce a concerted transcriptional decline at genes promoting DNA repair.

**Figure 5.**
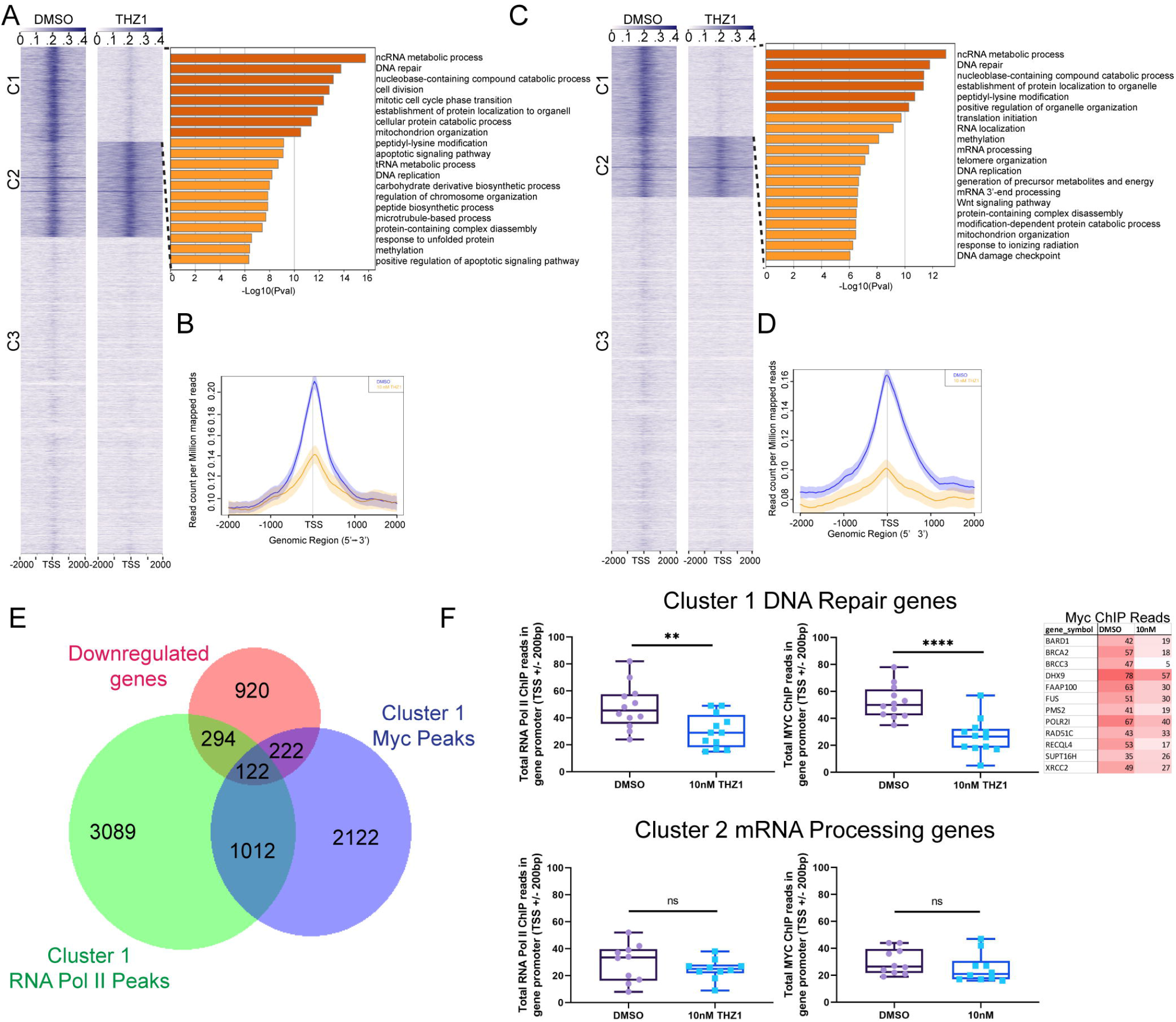
CDK7 Inhibition abrogates RNA Pol II and Myc promoter occupancy. (A) ChIP sequencing with RNA Pol II antibody performed on D458 cells treated with DMSO vs THZ1 10nM. Heatmap of normalized RNA Pol II occupancy at the TSS and, GO functional categories for cluster 1 genes effected by THZ1 treatment using metascape. Enrichment scores shown as −Log10(Pval). (B) Average read density of RNA Pol II ChIP-sequencing at the TSS. DMSO (blue) or 10nM THZ1 (orange). (C) ChIP sequencing with Myc antibody performed on D458 cells treated with DMSO vs THZ1 10nM. Heatmap of normalized Myc occupancy at the TSS and, GO functional categories for cluster 1 genes effected by THZ1 treatment using metascape. Enrichment scores shown as −Log10(Pval). (D) Average read density of Myc ChIP-sequencing at the TSS. DMSO (blue) or 10nM THZ1 (orange). (E) Venn diagram of gene overlap between cluster 1 of RNA Pol II ChIP, cluster 1 of Myc Chip, and genes downregulated by THZ1 treatment. (F) Box and whisker plots of the total number of reads within 200bp of the TSS for the top genes within the DNA Repair ontology of cluster 1 for RNA Pol II ChIP and Myc ChIP (top). Myc ChIP read numbers for top DNA repair genes, shown right. Color scale indicates high (red) to low (white) reads. Top genes within the mRNA processing ontology of cluster 2 for RNA Pol II and Myc ChIPs (bottom). Statistical analysis, two-tailed, unpaired t-test. ****, p<0.0001; **,p<0.01.

### Inhibition of CDK7 compromises homologous recombination and sensitizes MB cells to ionizing radiation

Transcriptional gene expression analysis revealed downregulation of genes mediating DNA repair and homologous recombination in addition to positive enrichment of genes responsible for apoptosis (Figure 6A, S5). Volcano plots of homologous recombination and DNA repair gene sets show individual genes involved in homologous recombination such as *Brca2*, *Rad51*, and *Rad50* are decreased upon THZ1 treatment (Figure 6B). Likewise, Gene Ontology analysis showed within the downregulated gene set the top genes effected were involved in DNA repair and homologous recombination (Figure 3F). To further validate if treatment with THZ1 reduced RNA Pol II occupancy at individual promoters of genes involved in homologous recombination, we examined the promoters of *Brca2*, *Rad51c*, a *Rad51* paralog, and *Rad51*. RNA Pol II promoter occupancy is noticeably depleted on *Brca2* and *Rad51c* promoters (Figure 6C). Examining the R*ad51* promoter, RNA Pol II is localized to a region 10kb upstream from the TSS demarcated by H3K27ac suggesting the presence of an enhancer. With THZ1 treatment RNA Pol II peaks are substantially reduced in this location (Figure S5). *Brca2* and *Rad51* are known targets of MYC, thus MYC occupancy was also examined. Correspondingly, MYC promoter peaks were reduced upon THZ1 treatment on the *Brca2* promoter, *Rad51c*, and *Rad51* promoters further validating our ChIP-sequencing results (Figure 6C and S5).

**Figure 6.**
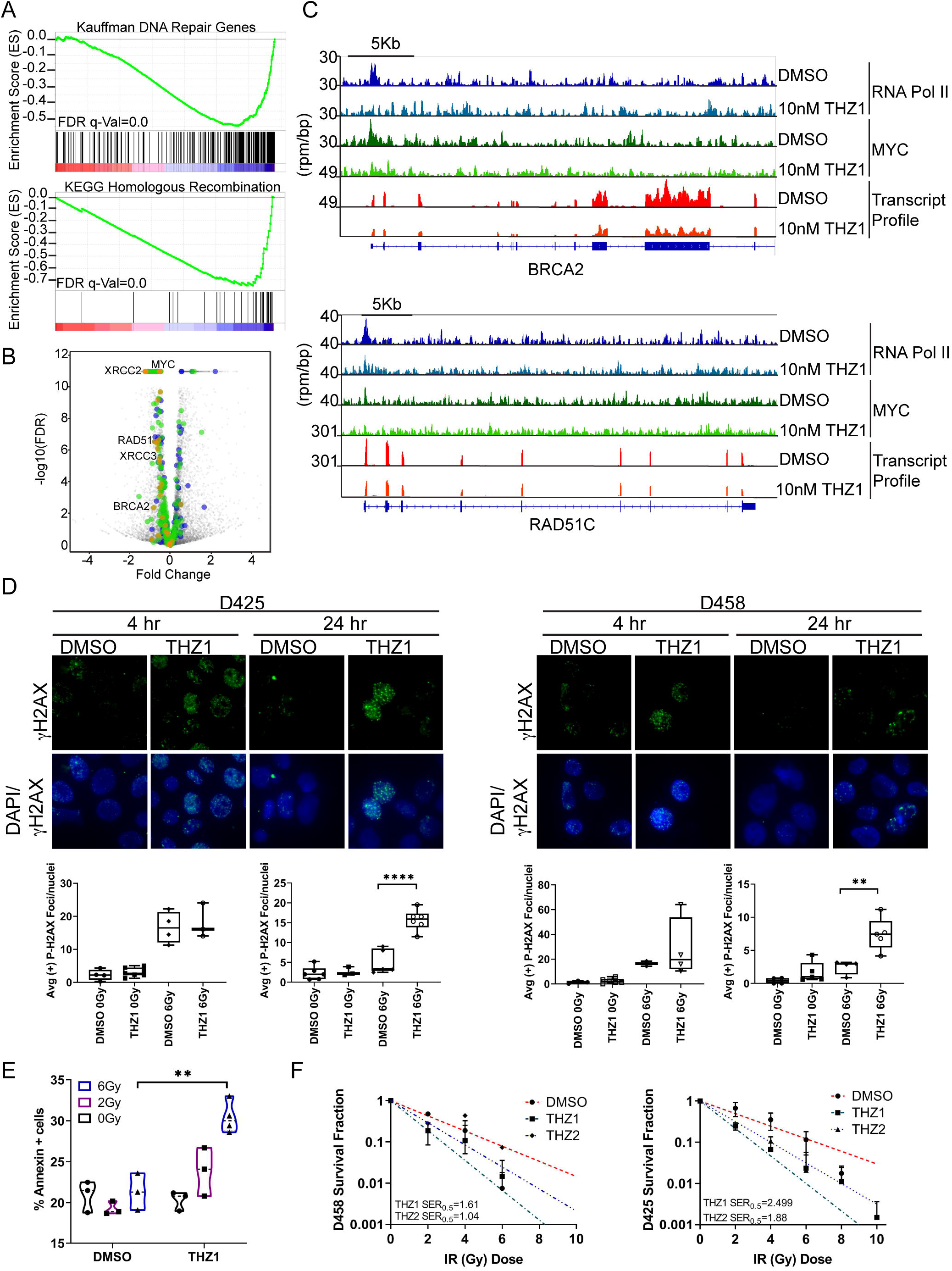
CDK7 inhibition minimizes DNA damage response enhancing susceptibility to ionizing radiation (A) GSEA from THZ1 D458 treatment RNA-seq. Kauffman DNA repair, KEGG Homologous Recombination. FDR q-Value=0.0. (B) Volcano plot of Kauffman DNA repair (green), KEGG homologous recombination (yellow), and MYC targets (blue). (C) Individual ChIP gene tracks of RNA Pol II, MYC signals, and RNA transcript profile for DMSO and THZ1 10nM D458 treatments. Y-axis signal density (rpm/bp) of BRCA2 promoter and RAD51C promoter. (D) Immunofluorescence of P-H2AX (green) shown at 100X and Dapi. D425 (left) and D458 (right) cells treated with THZ1 IC20 and exposed to 0 Gy or 6 Gy were fixed at 4hrs and 24hrs. Quantification of P-H2AX foci/nuclei (below). Statistical analysis, one-way Anova, ****,p<0.0001; **,p<0.001. (E) Annexin V (+) staining assay. D458 cells treated with DMSO or THZ1 200pM and ionizing radiation at 0, 2, and 6Gy were stained for Annexin V and measured by guava flow cytometery. Violin plot of the % (+) Annexin V stained cells (y-axis) is shown. Experiments performed in triplicate. Statistical analysis, two-way Anova. **,p<0.005. (F) Survival fraction plots of D458 (left) and D425 (right) from cells treated with DMSO or THZ1 200pM or THZ2 200pM and increasing ionizing radiation 0,2,4,6,8,10 Gy. Cells were grown in methylcellulose and colonies counted after 10 days. Survival fraction (y-axis) is plotted versus ionizing radiation (x-axis). Survival enhancement ratio is shown for THZ1 and THZ2.

The disruption of repair mechanisms would impair tumor cells defense mechanisms against commonly used therapeutic strategies such as ionizing radiation. We found that THZ1 treatment blocked RNA Pol II pausing at promoters of these specific genes, thus showing CDK7 chemical inhibition may sensitize MYC-amplified cells to DNA damage by preventing transcriptional amplification of DNA damage responders. To determine if MYC-amplified cells were more susceptible to ionizing radiation after THZ1 treatment, D458 and D425 cells were exposed to THZ1 IC_20_ and increasing dosage of radiation (0,2,4,6,8,10Gy). Initial immunofluorescence staining for P-γH2AX at 4hrs shows an accumulation of DNA DSBs in DMSO and THZ1 treated cells exposed to 6Gy compared to 0Gy for control (Figure 6D). However, at 24 hrs post radiation, THZ1 treated cells maintain a significantly higher level of DNA DSBs compared DMSO indicating a deficiency in DNA repair mechanisms (Figure 6D). Treatment with THZ2 replicated these results (Figure S5). To further assess the disruption of homologous recombination we performed immunofluorescence of RAD51 and RPA1 in D458 cells treated with IC_20_ THZ1 and irradiated at 4Gy. Consistent with decreasing HR function, a noticeable decrease in RPA1 and RAD51 foci was observed with THZ1 treatment (Figure S5).

We then examined induction of apoptosis by Annexin V staining of IR exposed and THZ1 treated D458 cells. Exposure to THZ1 along with 2 or 6Gy increased Annexin V positivity by >10% compared with DMSO suggesting increased sensitivity to IR with THZ1 treatment (Figure 6E, S5). The sensitivity enhancement ratio (SER) was then evaluated to determine the dosage of radiation with and without THZ1 or THZ2. A methylcellulose assay of the 0-10Gy IR exposed with THZ1 or THZ2 treated D458 and D425 cells were plated as before and monitored for growth after 10 days. In comparison to DMSO, D458 and D425 cells exposed to 20pM (10-fold less the IC_20_) THZ1 or THZ2 reduced the level of radiation needed to prevent independent colony growth. In D458 cells a 2Gy radiation dose results in a survival fraction of 0.47 and with THZ1 or THZ2 treatment the survival fraction is reduced to 0.28 (THZ2) and 0.18 (THZ1) (Figure 6F). Similarly, In D425 cells, survival after exposure to 2Gy radiation dose was 0.67 and with the addition of THZ1 treatment survival reduced to 0.24 (Figure 6F). In order to achieve similar decreases in survival without drug treatment the radiation dose would need to increase to 4Gy in both cell lines. At 50% survival the SER score for THZ1 of 1.61 and THZ2 of 1.04 in D458 and 2.49 and 1.88 in D425 signifies a synergism between THZ1 or THZ2 treatment and radiation exposure (Figure 6F).

### In vivo THZ2 treatment augments radiation sensitivity in group 3 MB cells

Given the observation that CDK7 inhibition sensitizes cells to ionizing radiation, we sought to evaluate whether the combination of THZ2 and ionizing radiation could be applied to the D458 orthotopic mouse model. Cerebellar injections of nude mice with D458 bioluminescent cells was subsequently followed by mouse randomization. Visible tumors were identified by IVIS imaging on day 11 post-injection (Figure 7A). Following tumor verification mice were irradiated at 1.5 Gy for 5 days and concurrent THZ2 treatment at 15mg/kg or vehicle was initiated. THZ2 intraperitoneal injections followed a similar timeline of 25 days. Tumors were tracked with bioluminescent imaging weekly which showed visible reduction in IVIS signals one week after radiation (Figure 7A). In the irradiated vehicle-treated cohort tumors recurred by week two, and at 65 days post injection 4 of 6 mice developed extensive tumor burden requiring euthanasia. THZ2 treated mice delayed tumor recurrence with little to no tumor signal up to three weeks after exposure (Figure 7A), with 5 of 6 mice still alive 65 days post injection. MRIs confirmed the presence of tumor growth in vehicle-treated cohort in comparison to THZ2 treated mice, where no tumors were detected for an additional 30 days beyond the last IVIS imaging (Figure 7B). The survival curve indicates a trend of survival enhancement in THZ2 treated mice where two mice failed to form tumors. While vehicle treated mice show faster tumor growth, one outlier did experience tumor deterioration after radiation which extended the life of the mouse. These results suggest THZ2 can extend the latency of tumor growth beyond treatment with radiation alone, and may be an effective treatment for combination therapy.

**Figure 7.**
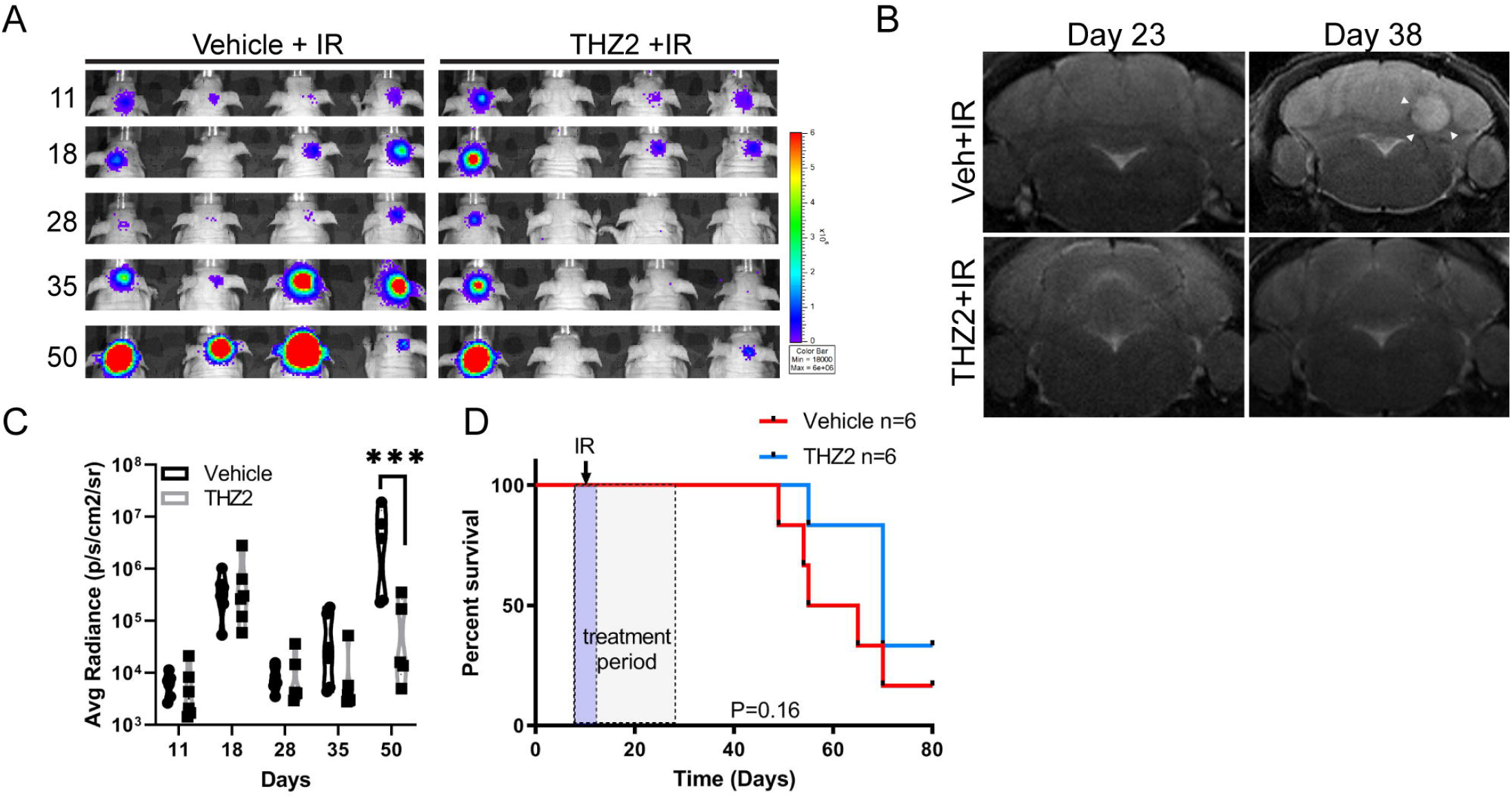
In vivo ionizing radiation with CDK7 inhibition. (A) Representative bioluminescence images of xenograft D458 vehicle (left) or THZ2 15mg/kg (right) treated with 1.5Gy over five days starting at day 15 post-injection. Color scales indicate bioluminescence radiance in photons/sec/cm^2^/steradian shown right. (B) Representative MRI of vehicle and THZ2 plus ionizing radiation treated D458 xenograft mice at 23 days and 49 days. White arrows indicate tumor. (C) Violin plot of mean total radiance in photons/sec/cm^2^/steradian. Statistical analysis, two-way Anova, ***,p<0.001. (D) Kaplan-Meier survival curve of D458 xenograft mice treated with vehicle or THZ2 and 1.5Gy for 5 days (violet box). THZ2 treatment period for 25 days indicated in shaded box. Statistical analysis log-rank (Mantel-Cox) test, p=0.16.

## Discussion

Group 3 medulloblastoma is largely devoid of germline driver mutations which can be therapeutically targeted, rather the subgroup population is most commonly linked with *Myc* amplification(Northcott et al., 2012; Ramaswamy et al., 2016). Amplification of *Myc* in medulloblastoma is associated with poorer clinical outcomes where tumors have a higher incidence of metastasis and recurrence(Northcott et al., 2012). These high-risk tumors are more aggressive in nature and less responsive to standard chemotherapy and radiation interventions. Standard radiation therapy for average-risk patients consists of whole brain and spine exposure, however high-risk patients still have no known optimal treatment where a combination of craniospinal irradiation with concurrent multiagent chemotherapeutic options is being explored(Northcott et al., 2019). Significant neurocognitive impairment predominates in high-risk patients and young children where experiences with long-term declines in IQ and memory are common due to ongoing nervous system development(Northcott et al., 2019; Ramaswamy et al., 2016). In a study that examined lowering the dosage of radiation to the posterior fossa the results showed worsening patient outcomes, thus radiation exposure remains static(Kann et al., 2016).

The genomic amplification of *Myc* causes widespread transcriptional dysregulation of specific gene clusters which aid in tumor cell propagation and evasion of cell death mechanisms(Bradner et al., 2017). In MYC-amplified tumors transcriptional amplification elevates replication stress inducing responses to DNA damage. However compensatory mechanisms sensed by ATR/CHK1 are also amplified by MYC alleviating cellular stress and enabling further tumor propagation(Campaner and Amati, 2012). This process imparts the ability to evade DNA damaging therapeutics and ionizing radiation leading to recurrent tumors and metastatic disease(Ryan et al., 2012). While uniquely targeting MYC amplification has remained problematic due to the absence of druggable catalytic domains, the potential to indirectly target MYC has shown promise (Bradner et al., 2017). Here we utilized a genome wide loss of function CRISPR-cas9 screen to identify specific targets in MYC-amplified medulloblastoma which have a pre-validated chemical inhibitor. The cyclin-dependent kinase CDK7 was found to mediate medulloblastoma growth in addition to protecting cells from apoptotic mechanisms.

CDK7 serves as a transcriptional regulator acting as the only catalytic component of TFIIH, phosphorylating the CTD of RNA Pol II on Ser5(Chen et al., 2018). The phosphorylation of RNA Pol II serves to position the RNA Pol II pre-initiation complex within the vicinity of the TSS priming RNA Pol II for the transition into progressive elongation(Chen et al., 2018; Core and Adelman, 2019). CDK7 mediates this transition by activating CDK9 within the P-TEFb complex, enabling RNA Pol II promoter proximal pausing and release into transcriptional elongation(Chen et al., 2018). While the systematic actions of TFIIH are absolutely necessary for global transcription, inhibition of the catalytic activity of CDK7 alone does not result in a genome-wide transcriptional blockade(Glover-Cutter et al., 2009; Kanin et al., 2007; Serizawa et al., 1993). Taking this into account, chemical inhibition of CDK7 has shown promise in multiple solid tumors which display hyper-active transcriptional dysregulation due to oncogene amplification(Christensen et al., 2014; Nagaraja et al., 2017; Wang et al., 2015). Evidence of CDK7 inhibition selectivity is found between TNBC which necessitates continuous active transcription compared to the ER/PR+ breast cancers that contain the traditionally targeted genetic alterations(Wang et al., 2015). CDK7 associated regulation of super-enhancers further defines the selectivity of CDK7 inhibition in transcriptionally dependent tumors (Chipumuro et al., 2014). Super-enhancers typically designated by an abundance of H3K27ac distal to the promoter stimulate oncogenic drivers. Super-enhancers are compelling new drug targets as they reveal epigenetic vulnerabilities that normal cells are not sensitive to. Thus, the identification of CDK7 in group 3 MB represents an opportunity to indirectly disrupt MYC transcriptional programs. Here, we find that CDK7 inactivation diminishes MYC and RNA Pol II promoter processivity on confined gene clusters containing DNA repair, DNA replication, and cell division networks.

Previous studies with THZ1 and CDK7 inhibition have found broad transcriptional deregulation in tumors with hyper-transcriptional amplification using concentrations higher than 100nM of THZ1. Off-target selectivity with larger concentrations contributed to the wide spread inhibition of gene expression. At the lower 10nM concentration we found a specific reduction of RNA Pol II and Myc at promoters of DNA damage regulatory networks including BRCA2 and RAD51. Concurrent with the ChIP data, our RNA-seq GSEA profiles show decreased enrichment of homologous recombination and DNA repair associated genes supporting the idea that treatment with THZ1 specifically deregulates medulloblastoma tumor cells defense mechanisms. Normally BRCA2 facilitates localization of RAD51 to double stranded breaks enabling connection of homologous DNA strands and adherence of breaks by DNA polymerization (Dietlein et al., 2014; Lord and Ashworth, 2007). Deregulation of either BRCA2 or RAD51 sensitizes cells to DNA damaging agents. We observed that decreased transcriptional amplification for *Brca2*, *Rad51c* and, *Rad51* impacted the level of ionizing radiation needed to induce cell death. Exposure to increased radiation and 10-fold lower dosage of THZ1 or THZ2 enhanced the sustained presence of DSBs indicating MB cells were more sensitive to radiation and less able to recover radiation induced DNA damage. Without CDK7 blockade cells are able to respond to the induction of DNA damage breaks in essence limiting the effectiveness of ionizing radiation on tumor cells. These data were further corroborated by Annexin V and SER data. Our collective SER data suggests that radiation exposure can be decreased significantly in combination with THZ1 or THZ2 treatment and achieve a comparable reduction in MB cell propagation. Additionally, the in vivo exposure of mice to radiation with THZ2 further informs these findings displaying a trend of delayed tumor regrowth and enhanced survival by 10 days.

The direct effect of CDK7 on homologous recombination is unclear, although, CDK7 activity effects both PLK1 and CDK1/2 which are direct regulators of BRCA2 and RAD51(Krajewska et al., 2015; Larochelle et al., 2007). Further, Myc is known to transcriptionally target several DNA repair genes including *Brca2* and *Rad51*. Treatment with THZ1 significantly deregulates RNA Pol II and MYC occupancy on isolated gene clusters compromising MYC transcriptional targets including *Myc* itself, DNA repair, ncRNA metabolism, and cell cycle. These actions suggest CDK7 inhibition represses RNA Pol II preinitiation complex assembly at the *Myc* promoter reducing transcription of *Myc* and its downstream targets. The following downregulation of homologous recombination genes may be further exacerbated by CDK7s’ activity as a CAK mediating CDK1/2, PLK1, and role in cell cycle. In support of this, another CDK7 inhibitor, YKL-5-124, was recently found to induce replication stress in SCLC cells independently of RNA Pol II CTD phosphorylation, blocking MCM2 at replication foci and inducing DNA damage and genomic instability (Zhang et al., 2020). These results further indicate additional deficits that could be exploited to target group 3 medulloblastoma.

As a whole CDK7 inactivation dismantles two mechanisms needed for medulloblastoma proliferation and survival. These data are particularly relevant given the recent development of clinical grade CDK7 inhibitors (Hu et al., 2019a; Hu et al., 2019b; Patel et al., 2018). CDK7 inhibition with SY-5609 has already entered clinical trials (NCT04247126). Our focus on DNA repair is relevant to increasing radiation therapy sensitivity in high-risk patients. Targeting DNA repair disturbs transcriptionally active group 3 medulloblastoma to enhance the impact of low dosage RT has potential to alleviate the neuronal deficiencies associated with treatment. Future developments of synergistic drug treatments with CDK7 inhibition are an alternative option to delaying intensive treatments, obviating the need of RT in children younger than 3 providing more unimpeded time for essential neuronal development.

## Methods

### Cell lines

D425 (RRID:CVCL_1275) and D458 (RRID:CVCL_1161) cell lines were kindly provided by Dr. Darell D. Bigner (Duke University Medical Center, NC). Cells cultured in DMEM supplemented with 10% FBS (Sigma-Aldrich), 1 mM sodium pyruvate (Gibco), 1× penicillin/streptomycin solution (Cellgro), and with 1× L-glutamine (Cellgro). ONS-76 (RRID:CVCL_1624) cells were obtained from Dr. James T. Rutka (The University of Toronto) and cultured in DMEM (Gibco, Carlsbad, CA) supplemented with 10% FBS (Sigma-Aldrich) and 1× penicillin/streptomycin solution (Cellgro). All cell lines were cultured at 37 °C with 95% air and 5% CO2. Cell lines were authenticated with STR fingerprinting using Globalfiler® System (Thermo Fisher Scientific) and processed on ABI 3500Xl Genetic Analyzer and mycoplasma testing was conducted with the Venor™ GeM Mycoplasma Detection Kit (Cat# MP0025-1kt, Sigma-Aldrich) both were done as recent as 1/9/2020. All cell lines were maintained for a maximum of 20 passages for the duration of experiments. Stable cell lines were established using gene specific mission shRNAs (Cat# TRCN0000000595, TRCN0000000592, TRCN0000230910) or non-targeting control (Sigma-Aldrich). Cells were plated at 5×10^5^ in a 6-well dish in growth medium containing 8μg/mL polybrene. Viral particles were added with 8μg/mL polybrene and incubated for 7 hrs. Transduced cells were disaggregated, grown, and selected with 1μg/mL puromycin for 14 days. Transformants were confirmed by qPCR and western blot.

### Compounds

THZ1 (Cat #9002215) was purchased from Cayman Chemical. THZ2 (Cat # HY-12280/CS-3245 was purchased from MedChem Express. Chemical compounds were diluted in DMSO at 10mM stock concentration.

### Annexin V Assay

Guava Nexin Assay #4500-0450 (Millipore). 2.0×10^5^ cells were collected and stained in 100 μl of Guava Nexin reagent for 20min then collected on Guava system flow cytometer (Millipore).

### CRISPR-Cas9 Screen

The CU Druggable Library (CUDL) consists of 8 gRNA per gene and targets 1095 genes curated from the 667 human genes targeted by 1194 FDA-approved drugs(Santos et al., 2017) and 428 genes encoding known drug targets involved in metabolism, protein modifications, signal transduction, and macromolecular transport biased for receptors and kinases(Wang et al., 2014) and druggable genes of interest from multiple research labs in University of Colorado’s Anschutz Medical Campus. The panel also consists of 500 non-targeting gRNA. The gRNAs were from http://www.broadinstitute.org/~timw/CRISPR; Cas-OFFinder tool was used to determine the potential off-targets(Bae et al., 2014) and Rule Set 2 score to measure the on-target activity of each gRNA(Doench et al., 2016). The oligo pool for the gRNA library was synthesized on a chip (CustomArray), PCR amplified and cloned into lentiCRISPRv2 (Addgene Plasmid #52961) and lentiGUIDE-puro (Addgene Plasmid #52963) (Sanjana et al., 2014) as described in(Joung et al., 2017; Wang et al., 2016) Next-generation sequencing of the amplified plasmid pool was performed to determine gRNA distribution.

### Methylcellulose assays

500 cells/3 mL were plated in a 1:1 mixture of 2.6% methylcellulose and complete growth medium. Cells were allowed to grow for ten days. Colonies were stained with nitrotetrazolium blue chloride (Sigma) at 1.5mg/mL in PBS for 24hrs at 37°C then counted.

### Neurosphere assay

D458 or D425 shNull and shCDK7 cells were serially diluted at 100,10, and single cell suspensions in neurosphere growth media (neurobasal medium, B-27+vitamin A, L-glutamine, pen/strep, EGF, and FGF). Cells were grown for fourteen days with media replacement every three days at 37°C. At fourteen days spheres were collected, disassociated, and replated at 100, 10, and single cell suspensions and allowed to grow for an additional fourteen days. Neurosphere proliferation and real time monitoring was done using the Incucyte^®^ S3 live cell imaging system (Essen Bioscience).

### RNA sequencing and Analysis

Total RNA was isolated in triplicate from D458 cells treated with 10nM THZ1 or DMSO using RNeasy Mini kit (Qiagen). RNA-seq was performed at Genomics and Microarray Core Facility, Anschutz Medical Campus using the illumina HiSEQ 4000 HT for 1×50 sequencing. High quality base calls at 95% ≥ Q30 were obtained with 53M-69M single reads. Statistical analysis of count data was performed with DESeq2 R package.(Love et al., 2014) Further analysis by GSEA was performed using the MSigDB (Mootha et al., 2003; Subramanian et al., 2005) and DAVID functional classification tool was used for gene ontology comparison (Huang et al., 2009a; Huang et al., 2009b).

### Western blot

Cells were lysed in RIPA buffer (Pierce, Thermo Fisher Scientific) containing an EDTA-free protease inhibitor (Roche), and protein concentrations were determined with the BCA Protein Assay Kit (Pierce, Thermo Fisher). 30μg of total protein was separated on a 4-20% gradient SDS-PAGE (BioRad). Primary antibodies α-RNA Pol II CTD S2 #13499, α-RNA Pol II CTD S5 #13523, α-CDK7 #2916, α-CDK9 #2316, α-PARP #9542, and α-cMYC #5605 (Cell Signaling Technology) were exposed overnight at 4°C. Secondary antibody, α-mouse-HRP Cat# 7076 and α-rabbit-HRP Cat# 7074 or α-Actin-HRP Cat# 12262 (Cell Signaling Technology), was exposed for 1hr at RT. Blots were developed with Luminata Forte Western HRP (Millipore) and imaged using Syngene GBox Chemi-SL1.4 gel doc. Western blots quantified using ImageJ.(Schneider et al., 2012)

### Chromatin immunoprecipitation

Cells were collected at confluency or approximately 1 million cells/IP. Cells were washed with 1xPBS, cross-linked with 1% paraformaldehyde in PBS at RT, and quenched with 2.5M glycine. Cells were then washed with ice cold PBS and lysed in 1mL RIPA buffer (150mM NaCl, 1% v/v Nonidet P-40, 0.5% w/v deoxycholate, 0.1% w/v SDS, 50mM Tris pH 8.0, 5mM EDTA) plus inhibitors (leupeptin 1μg/mL, aprotinin 1μg/mL, pepstatin 1μg/mL, benzamidine 1mM, and PMSF 1mM). Lysates were sonicated using Diagenode Bioruptor® with 25 cycles of 30 sec pulses and 90 sec intervals to shear DNA to ~500 bp fragments. Lysates were then cleared by centrifugation at 12800 rpm for 15 min at 4°C. 40 μl of protein A/G Sepharose beads (Millipore) were added and the lysate was pre-cleared for 1 hr at 4°C. 20 μl of protein A/G beads blocked with 1mg/mL BSA was mixed with the pre-cleared lysate, and immunoprecipitation was carried out by adding 5-10 μg of antibody (α-RNA Pol II #ab817 abcam, α-cMYC #13987, cell signaling technology) rotating overnight at 4°C. Beads were washed 2x with RIPA, 4X with IP wash buffer (100mM TrisHCL pH8.5, 500mM LiCl, 1%v/v Nonidet-P-40, 1% w/v deoxycholic acid), and 2x with TE buffer. Immunocomplexes were eluted at 65°C with elution buffer (70mM TrisHCl pH 8, 1mM EDTA, 1.5% w/v SDS). The eluate was brought to a final concentration of 200mM NaCl and reverse crosslinked at 65°C. The eluate was then treated with proteinase K, and DNA was isolated by phenol/chloroform extraction and ethanol precipitation. ChIP-DNA was quantified using the Qubit® dsDNA High Sensitivity Assay kit (Thermo Fisher).

### ChIP Sequencing and Analysis

Ovation UltraLow system V2 #0344 (Nugen) was used to prepare libraries as per protocol requirements. ChIP-seq libraries were sequenced on the Illumina Novaseq 6000 platform. 62M-69M reads with high quality base calls at 90%≥ Q30 were obtained. Bowtie2 was used to align the 150-bp paired-end sequencing reads to a reference human genome (hg19) downloaded from the UCSC Genome Browser. Unmapped and non-uniquely mapped reads were removed, and PCR duplicate reads were removed using samtools version 1.5. Peaks were called using MACS2 (v2.1.1.20160309) (Zhang et al., 2008) with default parameters. Peak locations were further annotated according to the known genes in hg38 and 3000◻bp of upstream and downstream of transcription start sites were considered as promoter regions using the R/Bioconductor package ChIPseeker (Yu et al., 2015).

### Bioluminescence Imaging

IVIS imaging was conducted on the Xenogen IVIS200 Bioluminescence system (Perkin Elmer). Mice were injected intraperitoneally with Luciferin #LUCK-5g, CAS 115144-35-9 (GoldBio) at 10μl/g body weight. Mice were anesthetized at 2% isoflurane/oxygen mixture then photons/counts were measured at 30 sec time intervals. Analysis of photon emissions was conducted using Living Image v2.60.1 software.

### Magnetic Resonance Imaging (MRI)

For in vivo MRI acquisitions, mice were anesthetized shortly before and during the MRI session using 1.5% of isoflurane/ oxygen mixture. Anesthetized mice were placed on a temperature-controlled mouse bed below a mouse head array coil and inserted into Bruker 9.4 Tesla BioSpec MR scanner (Bruker Medical, Billerica, MA). First, T2-weighted turboRARE images were acquired using the following parameters: repetition time (TR) = 3268 ms, echo time (TE) = 60 ms, RARE factor = 12, 8 averages, FOV = 20 mm, matrix size = 350×350, slice thickness = 700 μm, 24 sagittal and axial slices, in-plane spatial resolution = 51 μm. Then, diffusion weighted EPI sequence with 6 b-values was used using 4 axial slices covering the entire tumor lesions\ and unaffected brain tissue. Tumor regions were manually segmented on T2-weighted images by placing hand-drawing regions of interest (ROI) and the volume calculated as mm^3^. The apparent diffusion coefficients (ADC, s/mm^2^) were calculated from DWI maps as a criterion for tumor cellularity. All acquisitions and image analysis were performed using Bruker ParaVision NEO software.

### Irradiation of Culture Cells and Animals

Cultured cells were irradiated with a cesium-137 source (JL Shepherd Model 81-14R research irradiator) in 2 gray intervals from 0-10 gray. Animal radiotherapy was performed using the X-Rad SmART small animal irradiator (Precision X-Ray, North Branford CT). Under isoflurane anesthesia, mice received 7.5Gy to the cerebellum, in 5 fractions of 1.5Gy, delivered on consecutive days with pairs of 10mm-diameter lateral beams.

### Animal Experiments

For D458/D425 genetic KD and D458/D425/Med411 PDX treatment studies cells were collected and resuspended as a single cell suspension of 20000 cells/3 μL in serum free media. Intracranial injection of cells into outbred athymic Nude-Foxn1^nu^ mice (Jackson labs strain 07850) was done at 1.5mm lateral and 2mm posterior of lambda at 400 nanoliters/min. Mice were monitored for tumor growth daily and euthanized when 15% weight loss was reached. In vivo chemical treatment with THZ2, drug was diluted in 5% dextrose w/v and delivered via intraperitoneal injection at 15mg/kg for 25 days, daily. Vehicle was 10% DMSO diluted in 5% dextrose w/v. A Kaplan-Meier plot was used for survival calculations. Brain tissue was collected following euthanasia and fixed in 10% formalin.

### Histological and Immunofluorescence Staining and Microscopy

Fixed tissue was submitted to the University of Colorado Denver Tissue Histology Shared Resource for paraffin embedding, sectioning, and staining. Tissue was stained with α-CDK7 #2916, α-cleaved caspase 3 #9661, α-γH2AX #9718, α-RPA70/RPA1 #2267 (Cell Signaling Technology), α-Ki67#RM-9106 (Thermo Fisher) and α-RAD51 #NB100-148 (Novus Biological). Images were captured on an Olympus Bx43 light microscope and Keyence BZ-X700 series microscope at 4X, 20X, 100X then quantified with ImageJ.

### Study Approval

All animal procedures were performed in accordance with the National Research Council’s Guide for the Care and Use of Laboratory Animals and were approved by the University of Colorado Anschutz Medical Campus Institutional Animal Care and Use Committee.

### Statistical Analyses

All experiments were performed with at least three independent replications. All data was collected in Excel, and Graphpad Prism 8 statistical software was used for analysis. The mean ± SD is graphed. P-values <0.05 were considered significant, where p-value<0.05 (*), p-value<0.01(**), p-value<0.001 (***), p-value<0.0001 (****). Unpaired, two-tailed, T-tests were used for two-group comparisons, and ANOVA analysis and Dunnett’s test for multiple group comparisons.

## Supporting information

Full western images

## Declarations

### Consent for publication

Not applicable

### Availability of data and materials

The datasets used and/or analyzed during the current study are available on GEO database.

### Author Contributions

RV and BV designed the study and wrote the manuscript. BV conducted the experiments, data analysis, and prepared figures. KS, MJ, SKh, and SV created and SF performed CRISPR-cas9 screen (F1). ED performed ChIP-seq data processing and alignments (F4). AP performed mouse injections. DW performed RNA-seq analysis and GSEA (F3,6). NS performed MRI analysis (F4,7). SK performed animal radiotherapy (F7).

## Acknowledgements

We would like to thank Dr. Darell D. Bigner (Duke University) for generously providing the D458 cell line used in this study. The authors appreciate the contribution made by the University of Colorado Denver Tissue Histology Shared Resource, supported in part by the Cancer Center Support Grant (P30CA046934). The University of Colorado Cancer Center Functional Genomics core facility for lentiviral constructs and the Genomics and Microarray Shared Resource for their assistance with RNA-Sequencing and ChIP-Sequencing. The authors thank Jenna Steiner from the Colorado Animal Imaging Shared Resource (AISR) for acquiring all mouse MRI scans and Benjamin Van Court for animal radiography. The AISR is supported by the University of Colorado Cancer Center, the <u>P30CA046934 Cancer Center</u> and S10 OD023485 High-End Shared Instrumentation grants (NJS).

