## Supplementary material for "Transcriptional control of DNA repair networks by CDK7 regulates sensitivity to radiation in Myc-driven Medulloblastoma": Full western images

### Slide 1
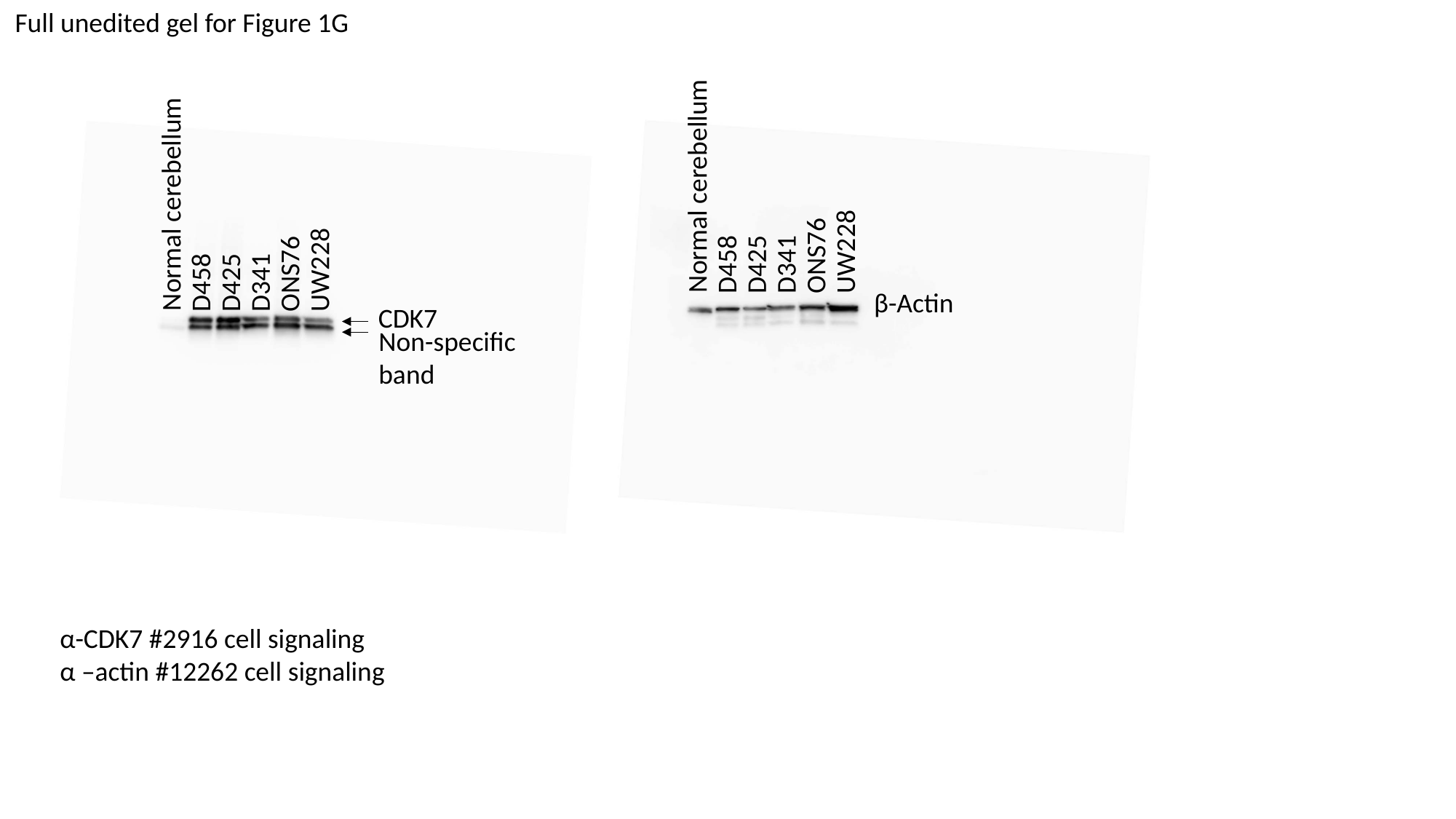

Full unedited gel for Figure 1G
Normal cerebellum
Normal cerebellum
UW228
ONS76
D458
D425
D341
UW228
ONS76
D458
D425
D341
β-Actin
CDK7
Non-specific
band
α-CDK7 #2916 cell signaling
α –actin #12262 cell signaling

### Slide 2
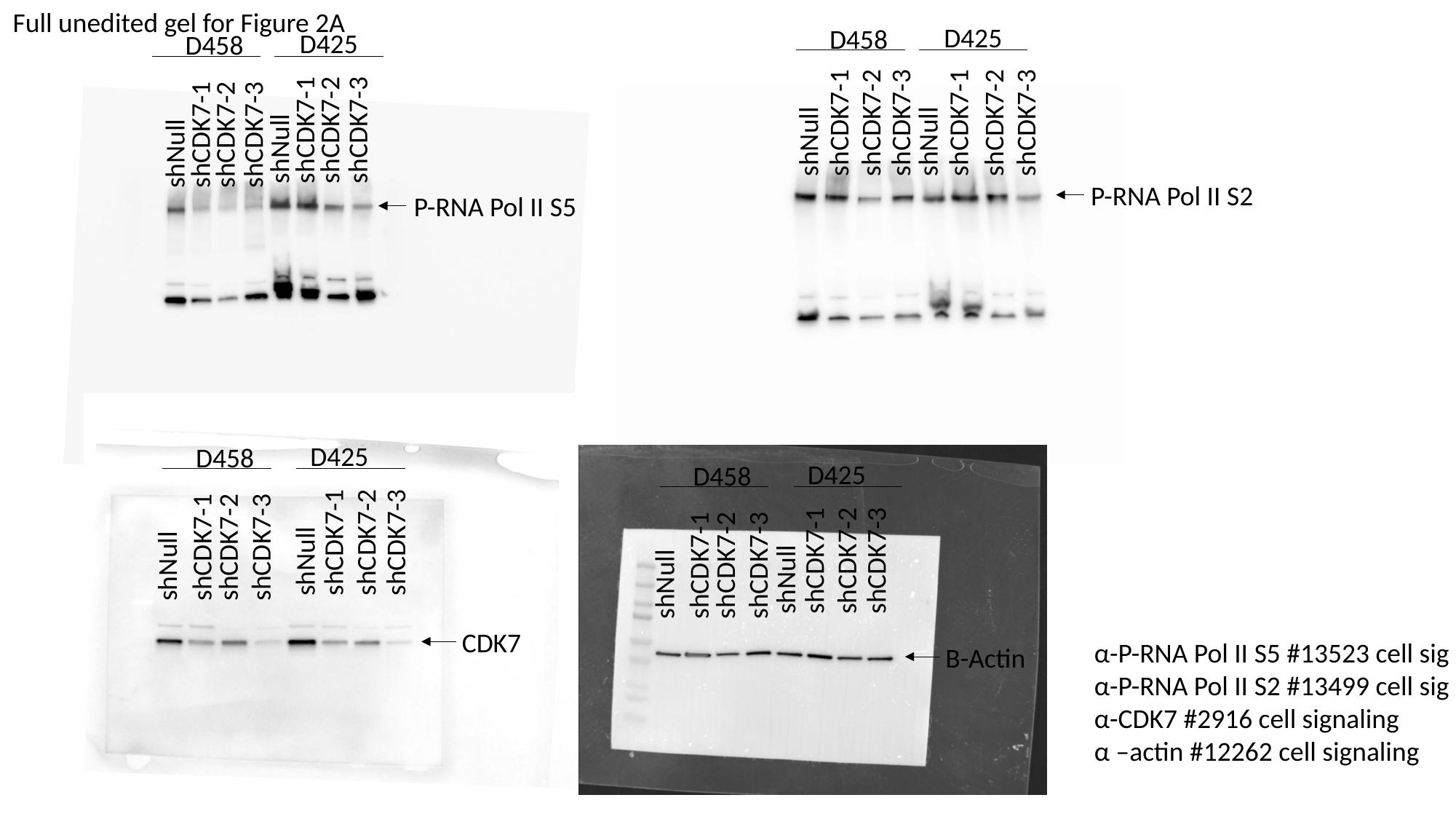

Full unedited gel for Figure 2A
D425
D458
D425
D458
shCDK7-1
shCDK7-2
shCDK7-3
shCDK7-1
shCDK7-2
shCDK7-3
shCDK7-1
shCDK7-2
shCDK7-3
shCDK7-1
shCDK7-2
shCDK7-3
shNull
shNull
shNull
shNull
P-RNA Pol II S2
P-RNA Pol II S5
D425
D458
D425
D458
shCDK7-1
shCDK7-2
shCDK7-3
shCDK7-1
shCDK7-2
shCDK7-3
shCDK7-1
shCDK7-2
shCDK7-3
shNull
shCDK7-1
shCDK7-2
shCDK7-3
shNull
shNull
shNull
CDK7
α-P-RNA Pol II S5 #13523 cell sig
α-P-RNA Pol II S2 #13499 cell sig
α-CDK7 #2916 cell signaling
α –actin #12262 cell signaling
Β-Actin

### Slide 3
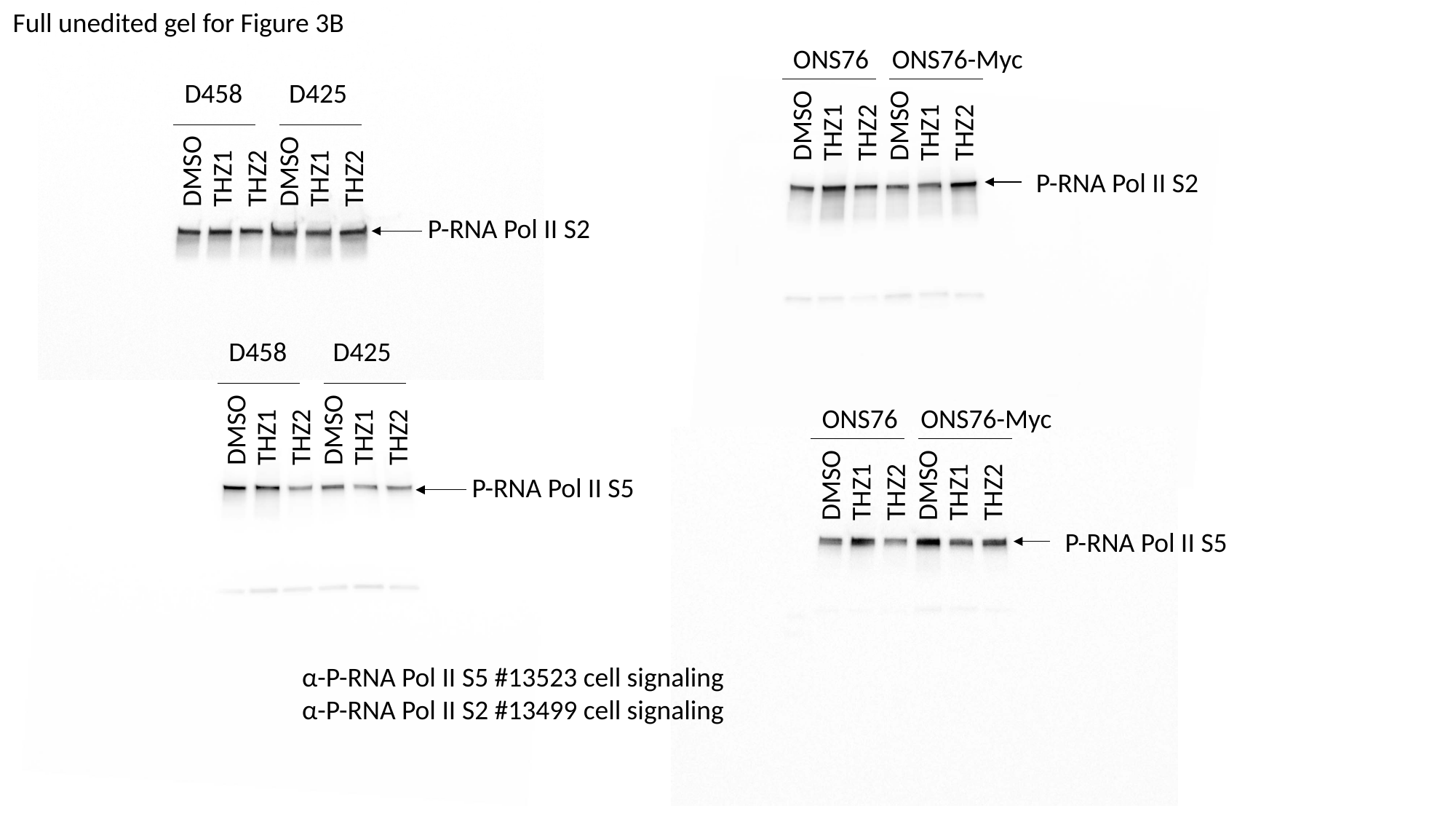

Full unedited gel for Figure 3B
ONS76
ONS76-Myc
D458
D425
DMSO
DMSO
THZ2
THZ2
THZ1
THZ1
DMSO
DMSO
THZ2
THZ2
THZ1
THZ1
P-RNA Pol II S2
P-RNA Pol II S2
D458
D425
ONS76
ONS76-Myc
DMSO
DMSO
THZ2
THZ2
THZ1
THZ1
DMSO
DMSO
P-RNA Pol II S5
THZ2
THZ2
THZ1
THZ1
P-RNA Pol II S5
α-P-RNA Pol II S5 #13523 cell signaling
α-P-RNA Pol II S2 #13499 cell signaling

### Slide 4
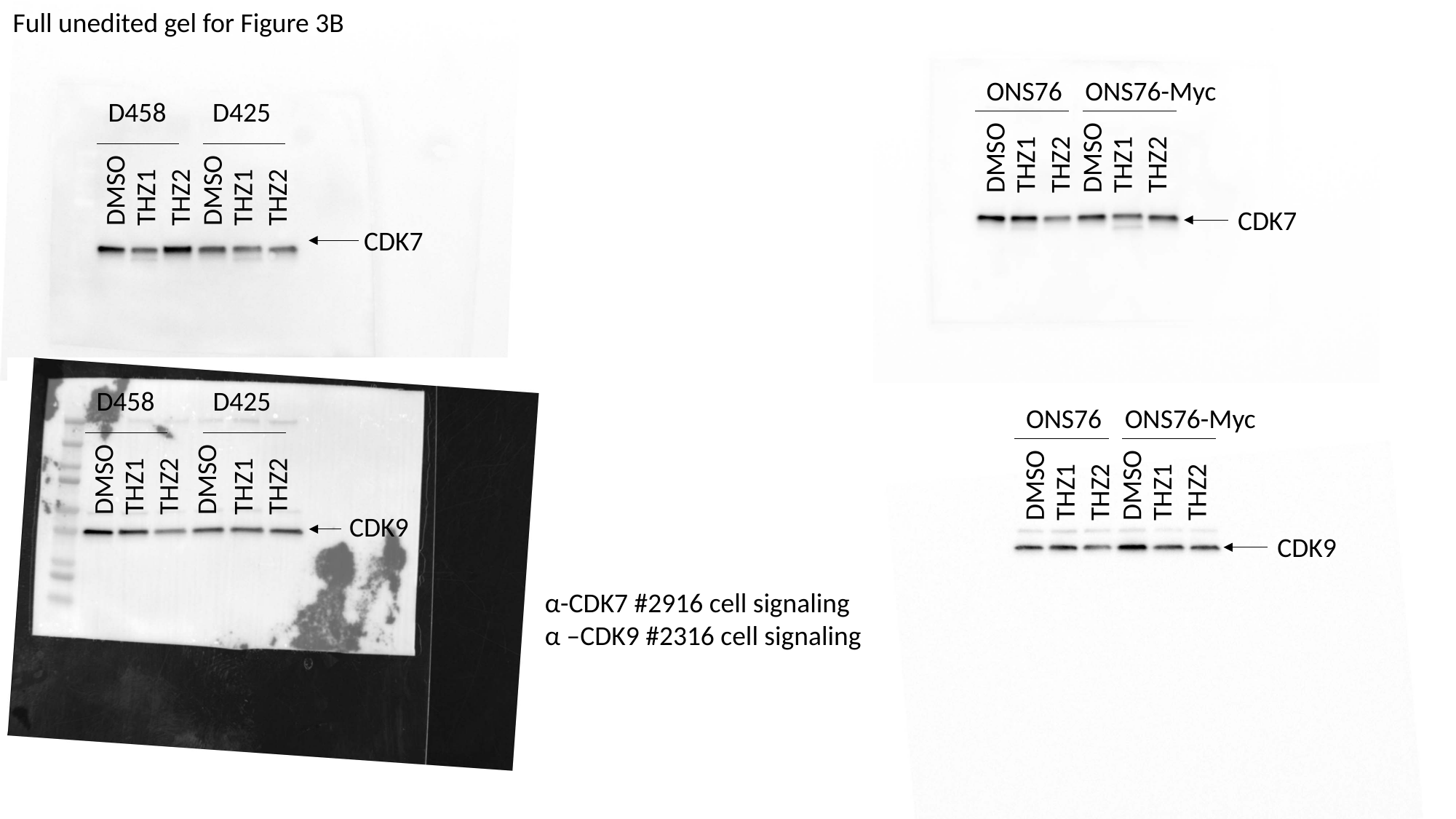

Full unedited gel for Figure 3B
ONS76
ONS76-Myc
D458
D425
DMSO
DMSO
THZ2
THZ2
THZ1
THZ1
DMSO
DMSO
THZ2
THZ2
THZ1
THZ1
CDK7
CDK7
D458
D425
ONS76
ONS76-Myc
DMSO
DMSO
DMSO
DMSO
THZ2
THZ2
THZ1
THZ1
THZ2
THZ2
THZ1
THZ1
CDK9
CDK9
α-CDK7 #2916 cell signaling
α –CDK9 #2316 cell signaling

### Slide 5
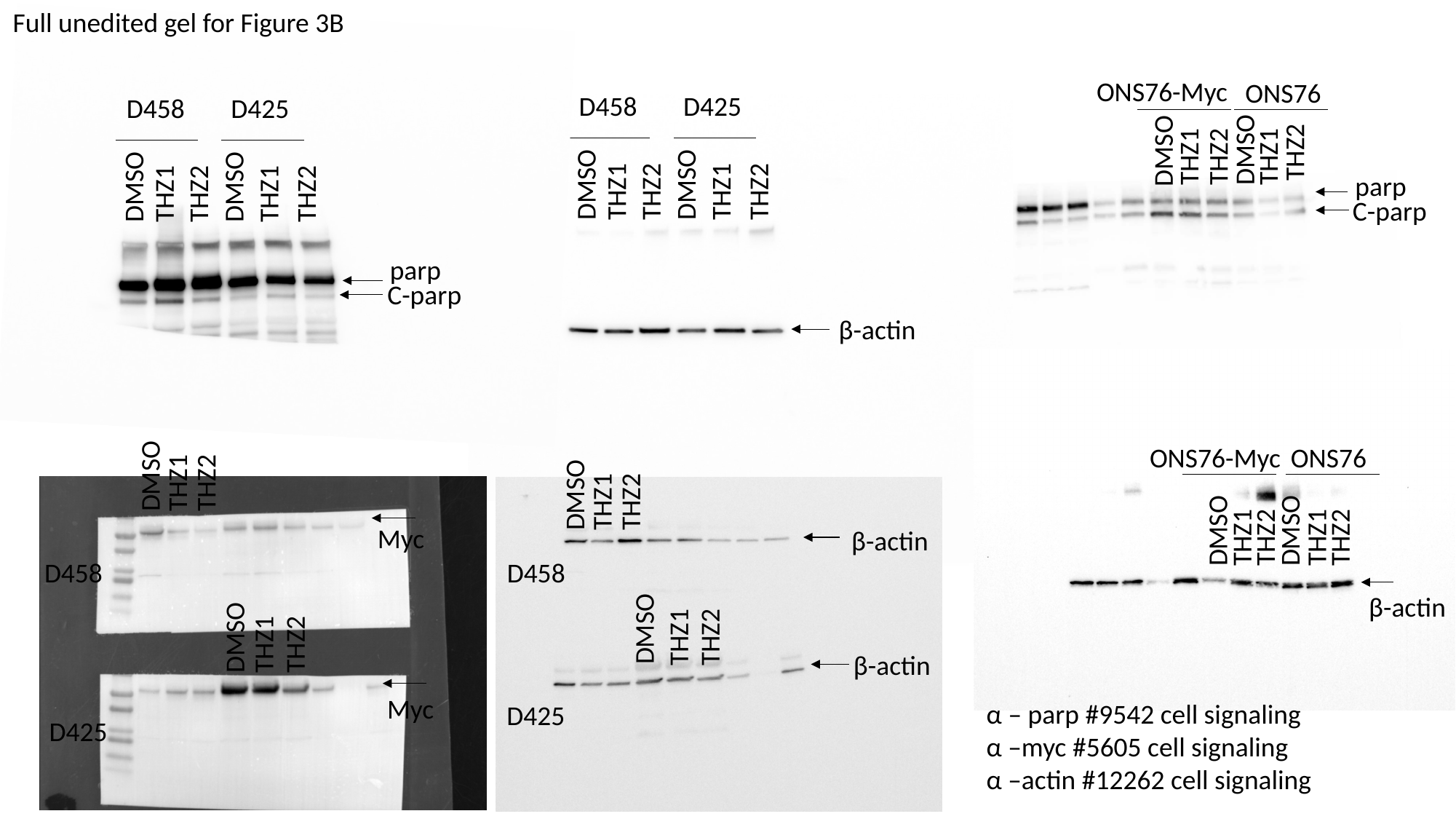

Full unedited gel for Figure 3B
ONS76-Myc
ONS76
D458
D425
D458
D425
DMSO
DMSO
THZ2
THZ1
THZ2
THZ1
DMSO
DMSO
parp
DMSO
DMSO
THZ2
THZ2
THZ1
THZ1
THZ2
THZ2
THZ1
THZ1
C-parp
parp
C-parp
β-actin
ONS76-Myc
ONS76
DMSO
THZ2
THZ1
DMSO
THZ2
THZ1
DMSO
DMSO
THZ1
THZ2
THZ1
THZ2
Myc
β-actin
D458
D458
β-actin
DMSO
THZ1
THZ2
DMSO
THZ2
THZ1
β-actin
Myc
α – parp #9542 cell signaling
α –myc #5605 cell signaling
α –actin #12262 cell signaling
D425
D425

### Slide 6
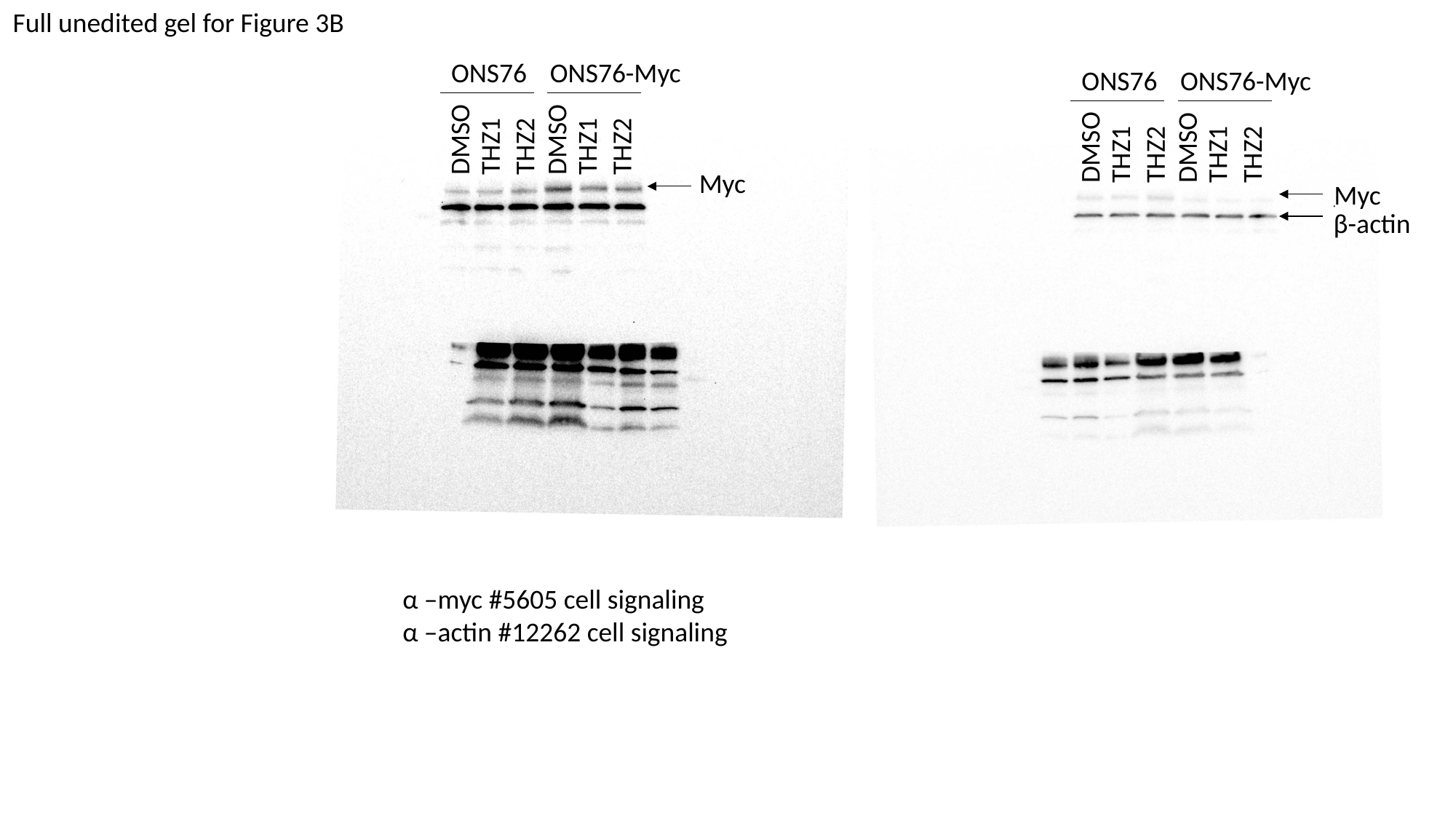

Full unedited gel for Figure 3B
ONS76
ONS76-Myc
ONS76
ONS76-Myc
DMSO
DMSO
THZ2
THZ2
THZ1
THZ1
DMSO
DMSO
THZ2
THZ2
THZ1
THZ1
Myc
Myc
β-actin
α –myc #5605 cell signaling
α –actin #12262 cell signaling
