## Supplemental Figures for "Transcriptional control of DNA repair networks by CDK7 regulates sensitivity to radiation in Myc-driven Medulloblastoma"

S1.

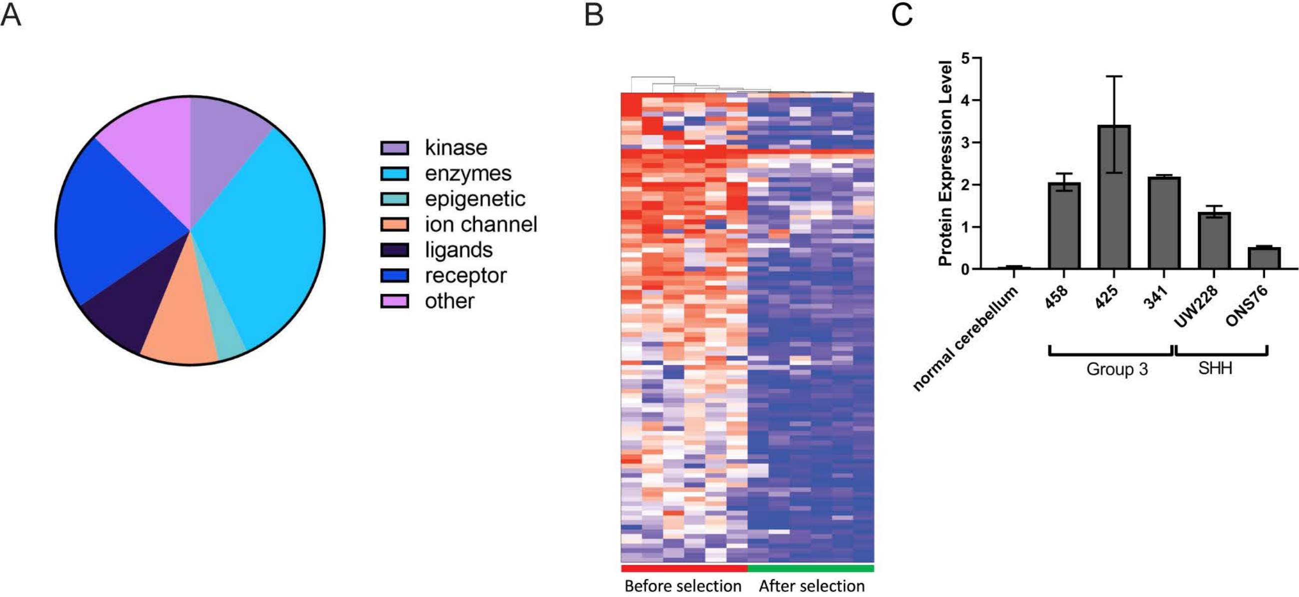

A
kinase
enzymes
epigenetic
ion channel
ligands
receptor
other
B
Before selection
After selection
C
Protein Expression Level
0
1
2
3
4
5
normal cerebellum
458
425
341
UW228
ONS76
Group 3
SHH

S2.

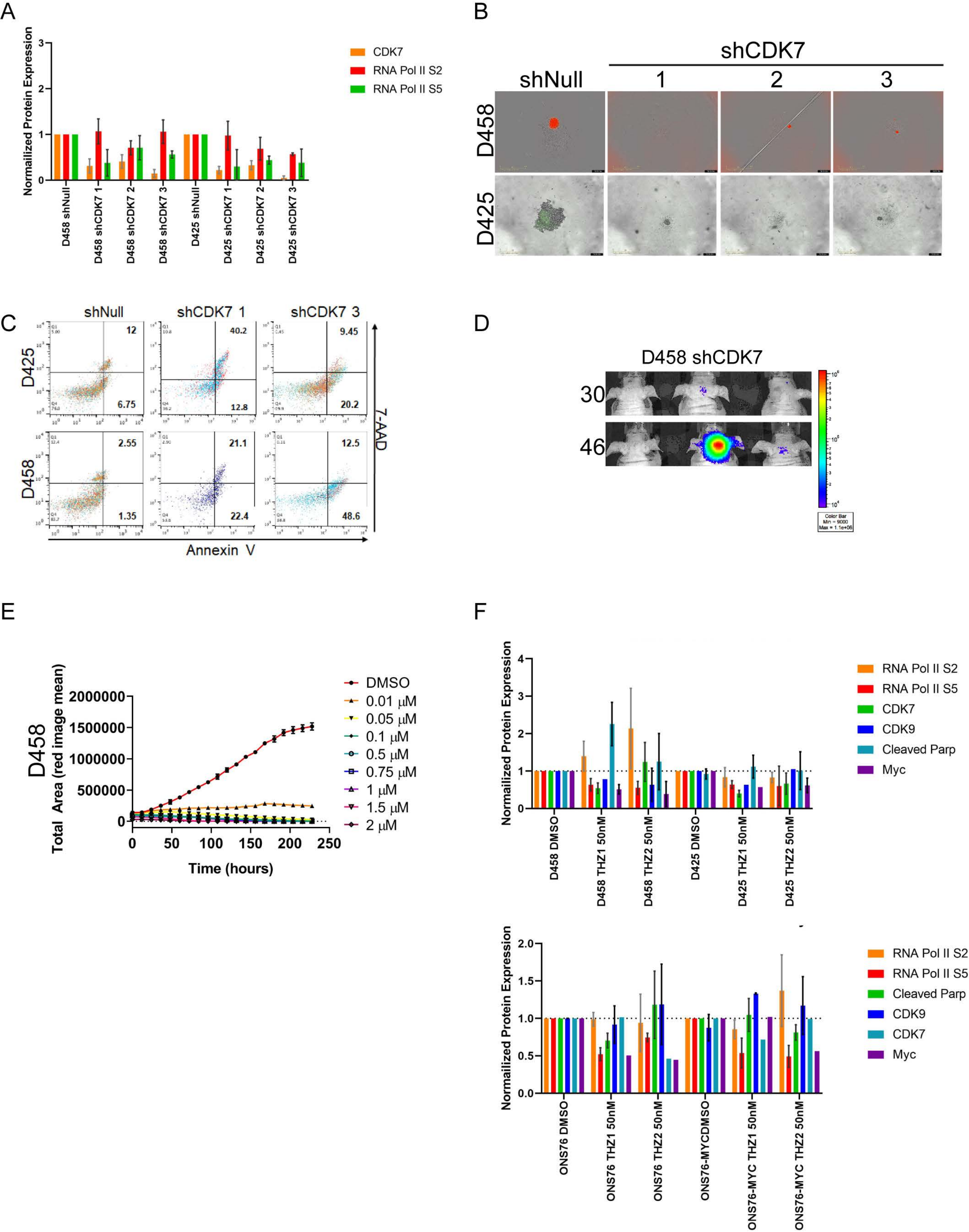
A
Normalized Protein Expression
CDK7
RNA Pol II S2
RNA Pol II S5
D458 shNull
D458 shCDK7 1
D458 shCDK7 2
D458 shCDK7 3
D425 shNull
D425 shCDK7 1
D425 shCDK7 2
D425 shCDK7 3
B
shCDK7
shNull
1
2
3
D458
D425
C
shNull
shCDK7 1
shCDK7 3
D425
D458
12
6.75
40.2
12.8
9.45
20.2
2.55
1.35
21.1
22.4
12.5
48.6
7-AAD
Annexin V
D
D458 shCDK7
30
46
Color Bar
Min = 9000
Max = 1.1e+06
E
D458
Total Area (red image mean)
Time (hours)
DMSO
0.01 µM
0.05 µM
0.1 µM
0.5 µM
0.75 µM
1 µM
1.5 µM
2 µM
F
Normalized Protein Expression
RNA Pol II S2
RNA Pol II S5
CDK7
CDK9
Cleaved Parp
Myc
D458 DMSO
D458 THZ1 50nM
D458 THZ2 50nM
D425 DMSO
D425 THZ1 50nM
D425 THZ2 50nM
Normalized Protein Expression
RNA Pol II S2
RNA Pol II S5
Cleaved Parp
CDK9
CDK7
Myc
ONS76 DMSO
ONS76 THZ1 50nM
ONS76 THZ2 50nM
ONS76-MYCDMSO
ONS76-MYC THZ1 50nM
ONS76-MYC THZ2 50nM

S3.

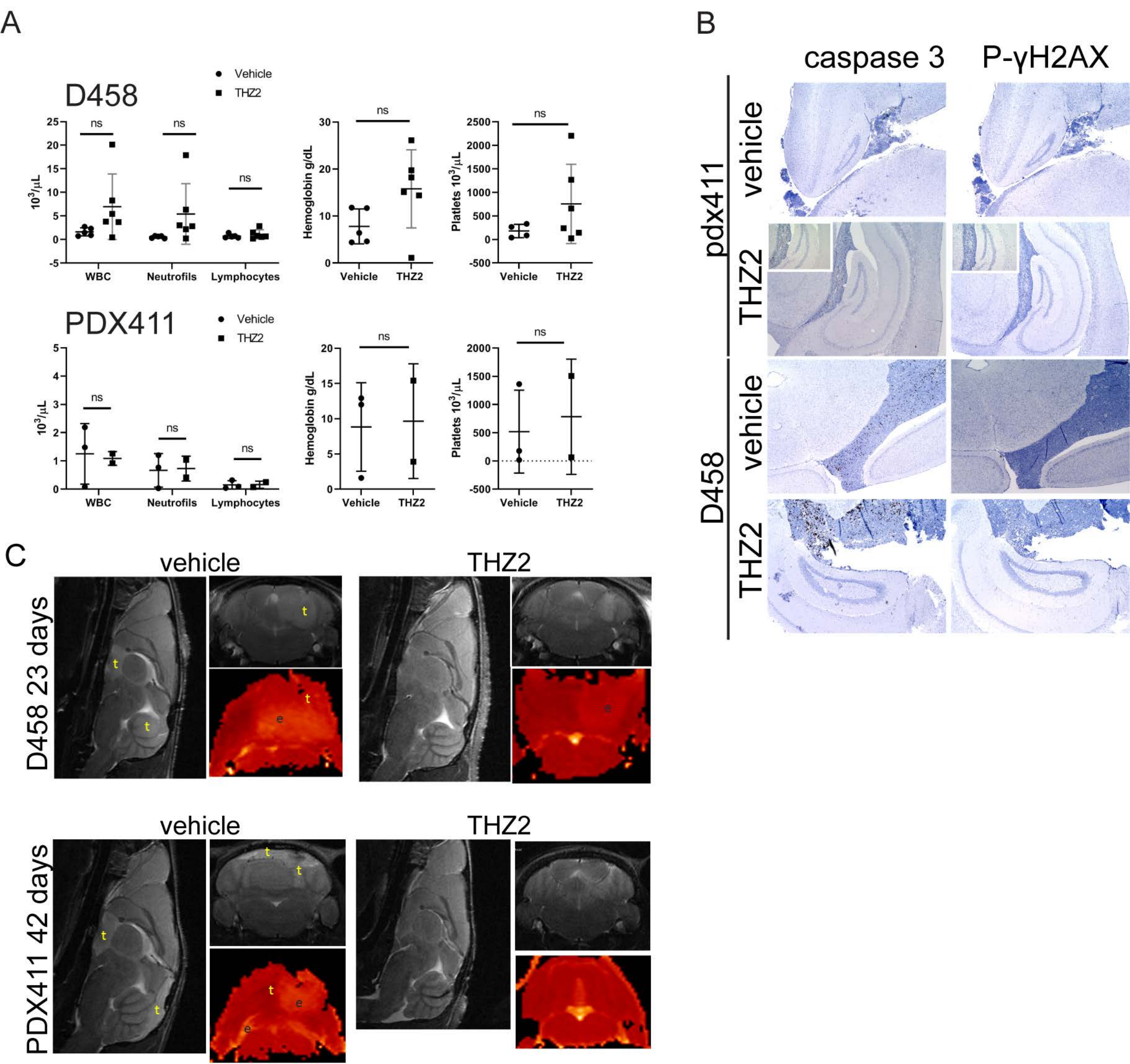

A
D458
Vehicle
THZ2
ns
$10^3/\mu L$
WBC
Neutrofils
Lymphocytes
Hemoglobin g/dL
Vehicle
THZ2
Platlets $10^3/\mu L$
PDX411
B
caspase 3
P-γH2AX
pdx411
vehicle
THZ2
D458
C
vehicle
THZ2
D458 23 days
PDX411 42 days
t
e

S4.

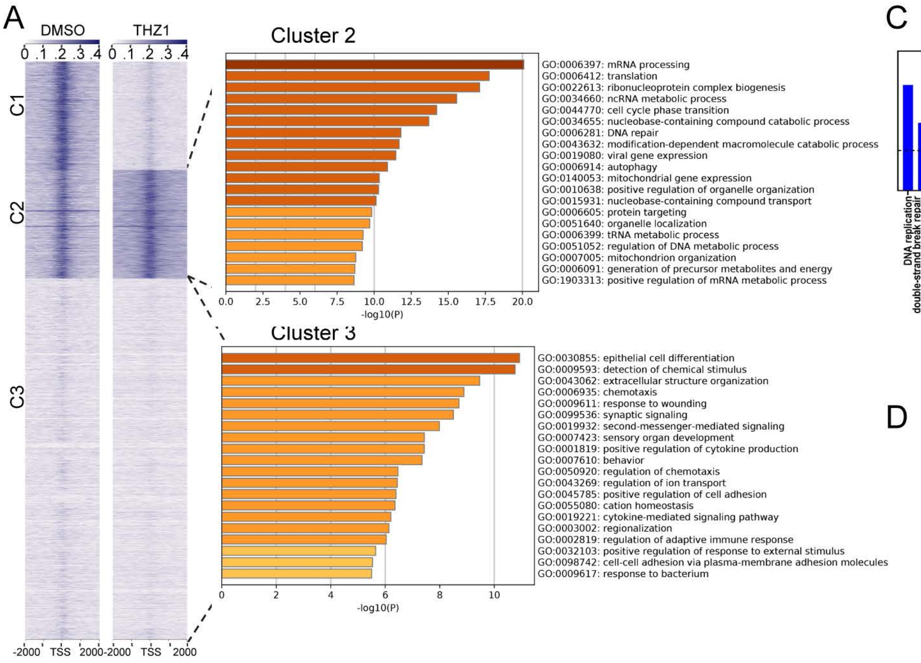


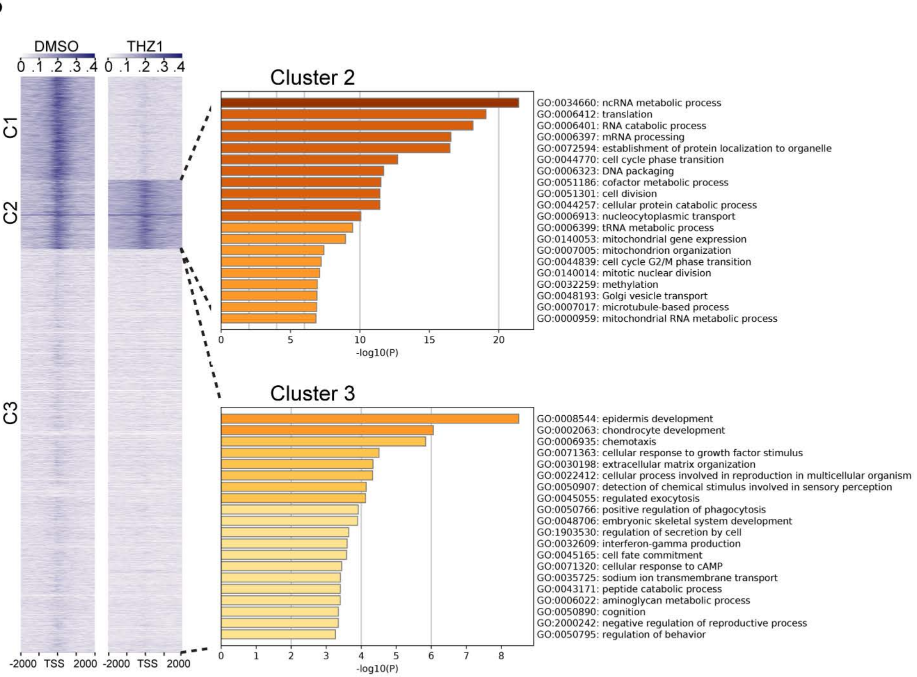


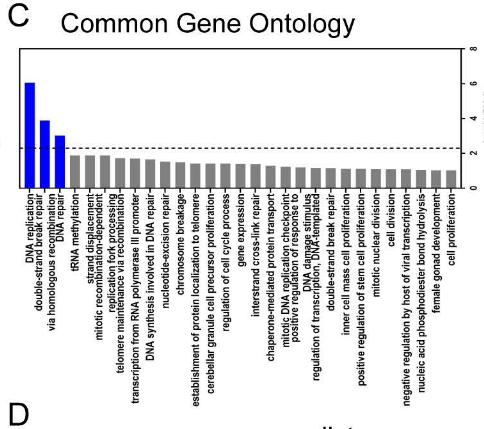


D

common gene list

| | | |
|---|---|---|
| ACADM | ISOC1 | SUPT16H |
| ACP2 | JAG2 | SUPT20H |
| ACTR5 | KIAA0196 | TARDBP |
| ANKRD31 | KIAA0895l | TELO2 |
| ANKS3 | LUC7L | TERT |
| AP3M2 | MAX | TIMM13 |
| ASL | MCM5 | TIMM8A |
| ATAD3A | METTL1 | TOMM40 |
| ATAD3B | METTL3 | TOP3A |
| ATF5 | MOCS3 | TPCN1 |
| ATG16L2 | MRPL36 | TRAK2 |
| ATP6V0C | MRPL46 | TRMT13 |
| ATPAF2 | MRPS11 | TTC30A |
| BCLAF1 | NABP1 | TTLL5 |
| BIVM | NADK2 | TXN |
| BOLA3 | NASP | UBE2G2 |
| BRCA2 | NDUFAF2 | UBP1 |
| C7orf50 | NHP2 | WDR90 |
| CCDC121 | NOL8 | XPO6 |
| CCDC138 | NOLC1 | ZBTB25 |
| CDC16 | NPDC1 | ZDHHC6 |
| CHRNA5 | NSUN2 | ZMPSTE2 |
| CLSPN | PA2G4 | ZNF12 |
| CTPS1 | PABPN1 | ZNF354B |
| DCAF4 | PEBP1 | ZNF558 |
| DDX18 | PEX10 | ZNF75A |
| DDX49 | PLCB3 | ZNF782 |
| DHODH | POLN | ZNF814 |
| DPF1 | POLR2F | ZNF839 |
| E4F1 | POLR3C | |
| EIF2B1 | PPP4R1 | |
| ENO1 | PRMT9 | |
| FAHD1 | PSMA3 | |
| FAM19A4 | PSMG1 | |
| FANCA | PUS1 | |
| FASTKD2 | RAD51 | |
| FBXW9 | RANBP10 | |
| FEN1 | RBM10 | |
| GINS4 | RBM25 | |
| GLDC | RBM6 | |
| GLYR1 | RCCD1 | |
| GPCPD1 | RELL2 | |
| GSS | RWDD2B | |
| GTF2H3 | SETD6 | |
| GTF3A | SETMAR | |
| HMGA2 | SFXN2 | |
| HSPB11 | SLMO1 | |
| HUS1 | SPATA13 | |
| IPO11 | SRSF10 | |

S5.

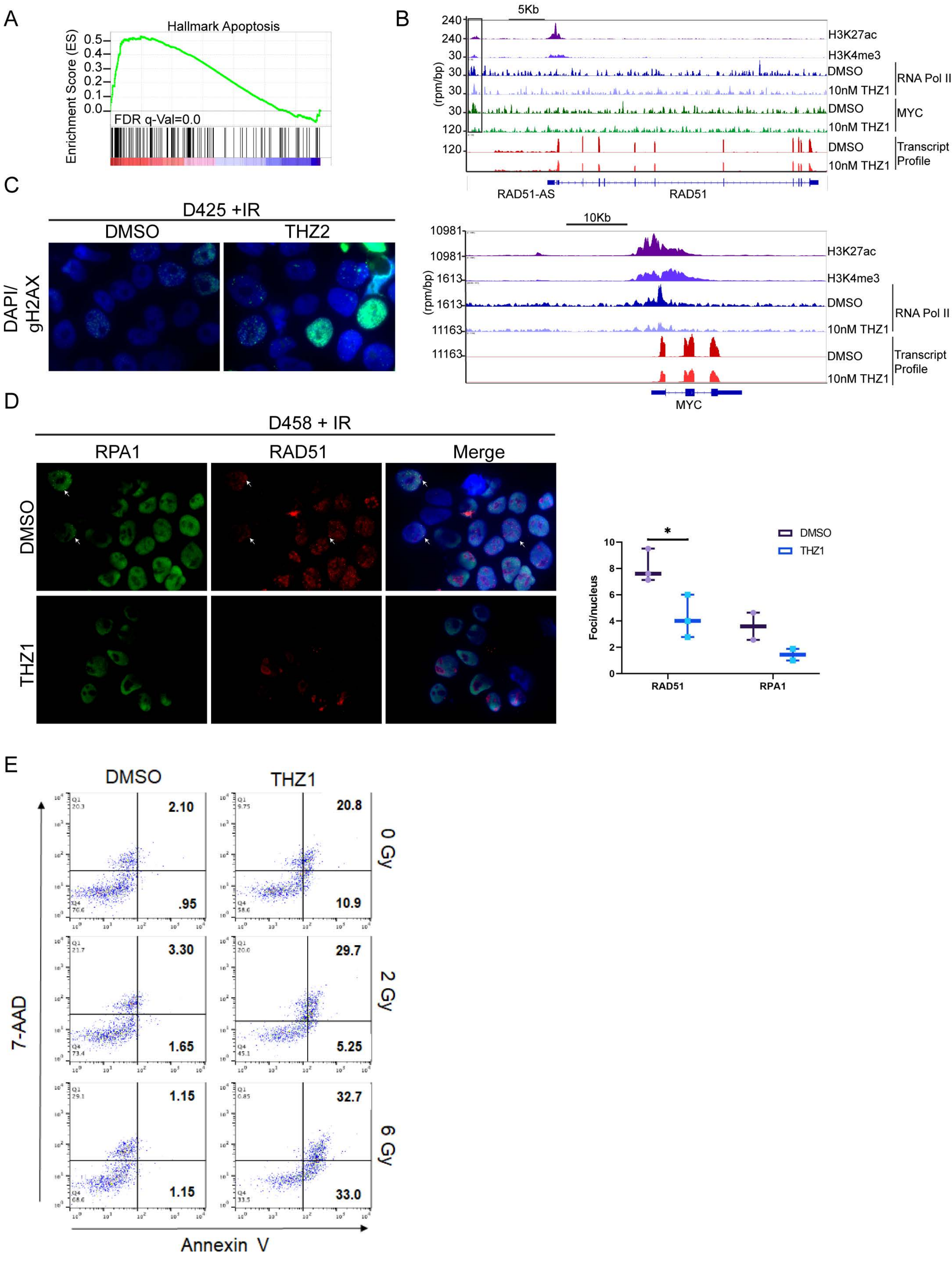
A
Hallmark Apoptosis
Enrichment Score (ES)
0.5
0.4
0.3
0.2
0.1
0.0
FDR q-Val=0.0
B
5Kb
(rpm/bp)
240
240
30
30
30
30
120
120
H3K27ac
H3K4me3
DMSO
10nM THZ1
RNA Pol II
DMSO
10nM THZ1
MYC
DMSO
10nM THZ1
Transcript Profile
RAD51-AS
RAD51
10Kb
10981
10981
1613
1613
11163
11163
H3K27ac
H3K4me3
DMSO
10nM THZ1
RNA Pol II
DMSO
10nM THZ1
Transcript Profile
MYC
C
D425 +IR
DMSO
THZ2
DAPI/ gH2AX
D
D458 + IR
RPA1
RAD51
Merge
DMSO
THZ1
DMSO
THZ1
Foci/nucleus
10
8
6
4
2
0
*
RAD51
RPA1
E
DMSO
THZ1
2.10
.95
20.8
10.9
0 Gy
3.30
1.65
29.7
5.25
2 Gy
1.15
1.15
32.7
33.0
6 Gy
7-AAD
Annexin V
