## Supplemental Figure legends for "Transcriptional control of DNA repair networks by CDK7 regulates sensitivity to radiation in Myc-driven Medulloblastoma"

Supplemental Figures.

S1. (A) Categorized drug panel pie chart. (B) Heatmap comparing before and after puromycin selection (C) Experiment performed in triplicate, data represents mean ± SD.

S2. (A) Quantified protein expression from protein immunoblot in Figure 2A. (B) Secondary neurospheres from D458 and D425 shCDK7 cells were plated at 10 cells/well and monitored for an additional 14 days. (C) Averaged Annexin V staining plot vs 7-AAD in D458 and D425 cells. (D) Representative bioluminescence images of xenograft D458 CDK7 knockdown or shNull Days post injection indicated on the left. Color scales indicate bioluminescence radiance in photons/sec/cm^2^/steradian (E) Proliferation of neurosphere D458 NucRed™ cells treated with increasing concentrations of THZ1. Cells were plated at 10 cells/well and monitored for growth on the Incucyte S3 system. Total average growth of neurospheres plotted by THZ1 concentration as measured by NucRed™ fluorescence. Experiments performed in triplicate, mean ±SD. (F) Quantified protein expression from protein immunoblot in Figure 3C.

S3. (A) Chemical toxicity as measured by WBC, neutrophils, and lymphocytes (10^3^/µl), hemoglobin (g/µl) and platelet counts (10^3^/µl) from D458 and PDX411 vehicle and THZ2 treated mice blood samples. (B) Representative immunohistochemistry staining for P-H2AX, and caspase 3 of D458 and PDX411 hippocampus from vehicle and THZ2 treated mice. Images taken at 4x and 20X.

S4. (A) ChIP sequencing with RNA Pol II antibody performed on D458 cells treated with DMSO vs THZ1 10nM. Heatmap of normalized RNA Pol II occupancy at the TSS and, GO functional categories for cluster 2 and cluster 3 genes effected by THZ1 treatment using metascape. Enrichment scores shown as -Log10(Pval). (B) ChIP sequencing with Myc antibody performed on D458 cells treated with DMSO vs THZ1 10nM. Heatmap of normalized Myc occupancy at the TSS and, GO functional categories for cluster 2 and cluster 3 genes effected by THZ1 treatment using metascape. Enrichment scores shown as -Log10(Pval). (C) (top) GO functional categories of common genes shown with enrichment scores as -Log10(Pval). Dashed line represents P< 0.005 with the most significant categories in blue. (bottom) Common gene list from venn diagram Figure 5E.

S5. (A) GSEA from THZ1 D458 treatment RNA-seq. Hallmark Apoptosis. FDR q-Value=0.0. (B) Individual ChIP gene tracks of H3K27ac, H3K4me3, RNA Pol II, MYC signals, and RNA transcript profile for DMSO and THZ1 10nM D458 treatments. Y-axis signal density (rpm/bp) of RAD51 promoter and Myc promoter. (C) Immunofluorescence of P-H2AX (green) shown at 100X and Dapi. D425 cells treated with THZ1 IC20 and exposed to 0 Gy or 6 Gy shown is 24hrs post radiation. (D) Immunofluorescence of RPA1 (green) and RAD51 (red) shown at 100X and Dapi. D458 cells treated with THZ1 IC20 and exposed to 4 Gy, shown is 24hrs post radiation. Quantification of foci/nucleus. Statistical analysis, one-way Anova, *,p<0.05. (E) Representative Annexin V staining plot vs 7-AAD in D458 treated with THZ1 and irradiated at 0, 2, 6gy.
